# Putative G-Quadruplex Structures in Cancer-Dysregulated Circulating lncRNAs and their G4-mediated Identification of Protein Interacting Partners

**DOI:** 10.64898/2026.05.27.728349

**Authors:** Deepshikha Singh, Arkadeep Ghosh, Shruti Mathur, Sanghati Patra, Safya Nasir, Rajesh K Hadiya, Bhaskar Datta

## Abstract

Circulating long non-coding RNAs (lncRNAs) have emerged as compelling cancer biomarkers. However, the structural features that mediate their extracellular stability and protein interactions remain largely unexplored. Here, we present the first systematic investigation of G-quadruplex (G4) motifs within cancer-dysregulated circulating lncRNAs and exploit these structures as molecular handles to identify associated RNA-binding protein (RBP) networks. From 283 circulating lncRNAs curated from the Lnc2Cancer 3.0 database, putative G-quadruplex-forming sequences (PQSs) were identified computationally using QGRS Mapper and G4Hunter, yielding four prioritized candidates — AGAP2-AS1, LINC00683, DLG1-AS1, and KRTAP5-AS1 — spanning 2G to 4G architectures. *In vitro* transcribed PQSs were validated for parallel G4 formation by circular dichroism spectroscopy, native polyacrylamide gel electrophoresis with thioflavin T staining, and reverse transcriptase stop assays, conducted under both standard buffer and simulated body fluid conditions to approximate the circulatory milieu. Electrophoretic mobility shift assays and isothermal titration calorimetry demonstrated nanomolar-affinity interactions between the G4-containing RNA and human serum albumin (HSA), the most abundant circulating protein. Cross-referencing G4-interacting proteins from the G4IPDB database with lncRNA-protein associations from LncTarD and NPInter, combined with RPISeq interaction predictions, identified ten candidate RBPs. A STRING-based protein-protein interaction (PPI) network was constructed at a confidence threshold of ≥0.7 and refined iteratively using experimental stability data to exclude proteins associated exclusively with the structurally weaker KRTAP5-AS1. The resulting network, centered on ELAVL1, IGF2BP1, hnRNPA2B1, and FUS, highlights a coordinated post-transcriptional regulatory module relevant to oncogenesis. This work establishes a novel, experimentally validated framework wherein G4 motifs serve as entry points for decoding the protein interactome of circulating lncRNAs, with implications for cancer diagnostics and RNA-targeted therapeutic strategies.

**Graphical Abstract:** 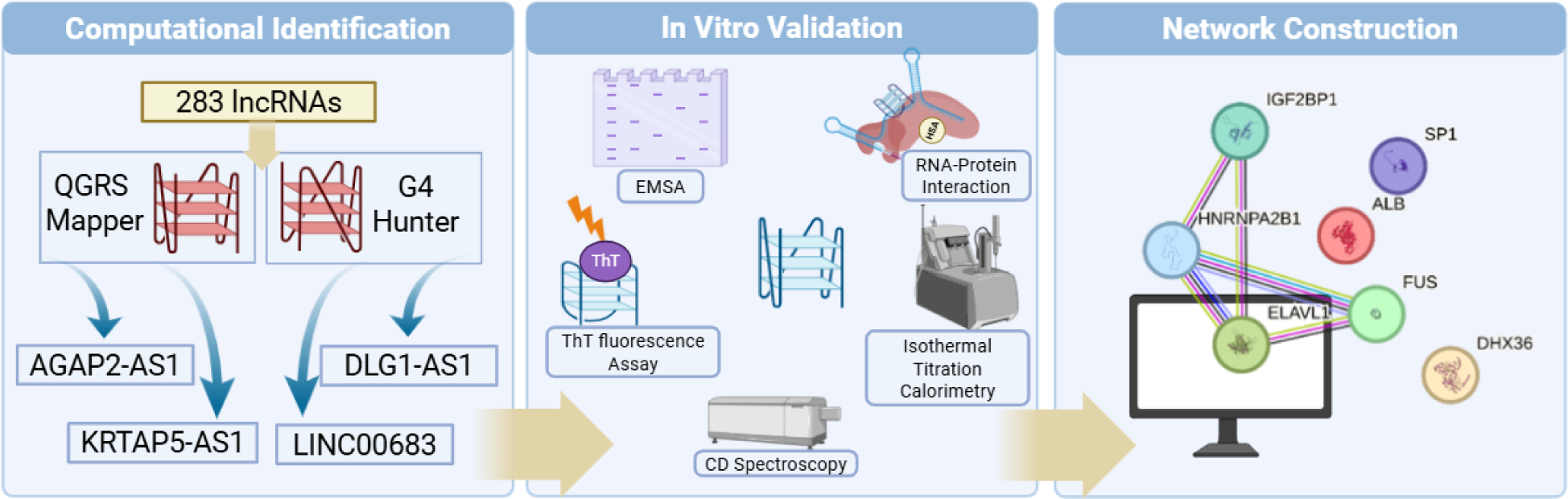

## 1. Introduction

Non-coding RNAs constitute the largest part of the transcriptome and have garnered significant interest over the past two decades towards understanding their physiological relevance broadly in terms of regulating gene expression *(Tassinari et al. 2021)*. Among non-coding RNAs, long non-coding RNAs (lncRNAs) are remarkable for their protein-like structure-function correlated activities. Dysregulated lncRNA levels signal disturbances in cellular regulation and are linked to many diseases, including diabetes, neurodegenerative and cardiovascular disorders, and various cancers *(Fenoglio et al. 2013)*. Circulating lncRNAs have emerged as important mediators and indicators of tumorigenesis, tumor growth and metastasis and advances in liquid biopsy enable minimally invasive detection and real-time monitoring. Despite challenges in validation and standardization, circulating lncRNAs show strong potential for diagnosis, prognosis, and therapy *(Beylerli et al. 2022)*. Circulating lncRNAs have been proposed as biomarkers for lung, breast, colorectal, gastric, hepatocellular, and prostate cancers *(Anfossi et al. 2018)*.

Prostate Cancer Antigen3 (PCA3), a lncRNA whose levels are highly upregulated in prostate cancer patients, is the first circulating lncRNA-based diagnostic test approved by the Food and Drug Administration (FDA), USA *(Day et al. 2011)*. Nandi Li et al. discovered the upregulation of plasma levels of HOTAIR in non-small cell lung cancer (NSCLC) patients *(Li et al. 2017)*. In another study, the levels of three lncRNAs, namely LINC01627, LINC01628, and ERICH1-AS1, detected in the plasma of lung cancer patients were found to be upregulated as compared to healthy controls *(Tang et al. 2015b)*. The lncRNA RP11-445H22.4 has been proposed as a diagnostic biomarker in breast cancer *(Xu et al. 2015)*. Graham et al. showed that 87% of colorectal cancer patients had upregulated levels of CRNDE in their plasma samples as compared to healthy individuals. Wang et al. identified a 3-lncRNA signature marker comprising SNHG-16, TUSC7, and RPL34-AS1 with diagnostic potential *(Wang et al. 2016)*. In hepatocellular carcinoma, Tang et al. reported that plasma levels of two lncRNAs, XLOC_014172 and LOC149086, were indicative of metastatic spread in diseased versus healthy controls *(Tang et al. 2015a)*. GIHCG lncRNA was found to be a prognostic biomarker in renal cell carcinoma, with serum levels directly correlated with renal carcinoma tissue levels *(He et al. 2018)*.

Several studies have shown that lncRNAs function through coordinated interactions with multiple RNA Binding Proteins (RBPs) rather than isolated one-to-one associations, making protein-protein interaction (PPI) network construction an important systems-level approach for understanding their biological roles. Studies have produced PPI networks through STRING and Cytoscape to analyse and identify functional hubs along with their regulatory modules from lncRNA-associated proteins in cancers like lung, colorectal, gastric, and hepatocellular carcinoma to reveal key pathways involved in RNA metabolism, splicing, proliferation, and metastasis *(Wu et al. 2016; Lan et al. 2020; Wang et al. 2022)*. Core RBPs such as ELAVL1, IGF2BP1, FUS, hnRNPA2B1, and hnRNPK are well-established participants in lncRNA-mediated regulation, often functioning together in transcript stabilization, phase separation, and post-transcriptional control *(Wippel et al. 2021)*. Similarly, DHX36 has been specifically implicated in resolving RNA G-quadruplexes and interacting with G4-containing lncRNAs, supporting its inclusion in our analysis *(Tippana et al. 2019)*.

The dominant approach for assessing the potential of lncRNAs as potential biomarkers of cancer is based on the comparison of healthy versus disease samples. Such studies often include correlation of lncRNA expression after treatment to assess robustness and responsiveness of the prospective biomarkers*(Senousy et al. 2021; Zhang et al. 2024)*. The stability of lncRNAs in the extracellular space stems from their encapsulation by extracellular vesicles or the presence of distinct secondary structures that can provide protection from nucleases. The provenance of lncRNAs in extracellular fluids has been linked to their association with proteins in a complex. It is plausible that prior to secretion, these lncRNAs might be binding to RBPs within a tumor cell thereby driving tumorigenesis *(Badowski et al. 2022)*. LncRNAs in the bloodstream may interact with suitable proteins present in the circulating system, such as albumin, which might aid in its stabilization in the bloodstream. We have previously sought to identify G-quadruplex (G4) motifs on cancer-dysregulated lncRNAs as a prospective structural anchor for inferring interacting proteins and as a target that can be leveraged for pharmacomodulation. To the best of our knowledge, circulating lncRNAs have yet to be systematically investigated regarding their ability to bear G4s. There is a dearth of studies on structural features of circulating lncRNAs that mediate interactions and contribute to extracellular stability. To date, no study has systematically interrogated RNA G4 motifs within circulating lncRNAs as structural determinants of protein partner recruitment

Human serum albumin (HSA) is the most abundant plasma protein and contains multiple positively charged and hydrophobic surface regions that enable moderate-affinity binding to RNA species, as reported for several microRNAs *(Botti et al. 2022a, 2022b)*. Collectively, these properties position HSA as a physiologically relevant partner *(Gerasimova et al. 2010)*. Serum albumin carries prognostic significance, and lower levels of HSA have been correlated with worse survival outcomes in colorectal cancer patients *(Boonpipattanapong and Chewatanakornkul 2006)*. Interestingly, the concept of albumin – a non-classical, non-RBP plasma protein – engaging G4s specifically was absent from literature prior to 2024. We have recently reported the strong association between HSA and G4s formed by ovarian cancer-dysregulated lncRNAs *(Singh et al. 2025)*. In circulating lncRNAs, G4 motifs could serve as useful entry points for identifying associated protein partners and for deriving broader PPI networks linked to their extracellular functions.

In this work, we envisage G4 motifs as structural hooks to investigate how circulating lncRNAs engage with proteins in their environment. By leveraging the unique capacity of G4 structures to recruit G4-recognising and RNA-binding proteins, we use G4-forming potential as a criterion to systematically prioritize circulating cancer-associated lncRNAs. Based on circulating lncRNA datasets, we have identified putative G-quadruplex-forming sequences (PQS), integrated sequence-based interaction predictions, and tried to map lncRNA-protein associations within cancer-relevant pathways. Experimental assessments of G4 interactions with HSA reveal high-affinity interactions that are 3 – 4 orders of magnitude stronger than prior RNA – HSA affinity measurements *(Botti et al. 2022a, 2022b)*. Further, we have refined the PPI network based on feedback from the in vitro experimental data. This work demonstrates a viable roadmap for exploiting G4 motifs as discovery handles to connect circulating lncRNAs to their interacting protein partners in circulatory fluid, as well as to illuminate their potential mechanistic contributions in cancer biology.

## 2. Materials and Methods

### 2.1 In silico identification of putative G-quadruplex-forming sequences in circulating lncRNAs

Circulating lncRNAs dysregulated in various cancers were retrieved from the Lnc2Cancer 3.0 database. The dataset was downloaded, and sequences for each lncRNA and transcript variant were obtained from the NCBI-Nucleotide database, filtering for Homo sapiens, linear ncRNAs, and validated or reviewed RefSeq status. Aliases were cross-checked using GeneCards, and sequences were compiled in FASTA format. PQS were identified using QGRS Mapper and G4Hunter. QGRS Mapper parameters were set to analyze PQS up to 45 nucleotides, with loop sizes of 0-36. G4Hunter analysis was performed using a sliding window of 45 and thresholds of 0.9 and 1.4. lncRNAs with PQSs showing high G-scores in both analyses were prioritized. Four lncRNA sequences with strong PQS overlap in QGRS Mapper and G4Hunter were selected for further *in vitro* experimental validation, including one 2G (KRTAP5-AS1), two 3G (AGAP2-AS1, LINC00683), and one 4G sequence (DLG1-AS1). The putative quadruplex-forming sequences identified within the selected circulating lncRNAs are hereafter referred to as PQS in the in vitro experimental protocols and results discussed in this work. For simplicity, we have used the name of specific lncRNAs interchangeably with their cognate PQS. Further, considering our objective of using G4 motifs as anchors for accessing lncRNA-binding proteins, we refer to the lncRNAs by their names in the in silico component of the work performed.

### 2.2 Identification and interaction prediction of lncRNA-Protein Partners

The identification of protein partners for cancer-associated circulating lncRNAs was carried out using the LncTarD 2.0 and NPInter v5.0 databases *(Zhao et al. 2019; Zheng et al. 2023)*. Database queries were performed for each lncRNA to annotate both direct and indirect RNA-protein interactions. Protein partners with the strongest evidence for interaction and disease relevance were prioritized for further analysis and functional enrichment studies. Subsequent to initial partner identification, interaction prediction between lncRNAs and candidate proteins was performed in silico using RPISeq, which employs both Random Forest (RF) and Support Vector Machine (SVM) classifiers to estimate binding probabilities from primary sequences *(Muppirala et al. 2011)*. Sequence data for lncRNAs and proteins were provided in FASTA format, and interaction scores above 0.5 for any one of the classifiers were considered high confidence. Pairs meeting this threshold were retained as candidates for exploring direct lncRNA-protein interactions.

### 2.3 Protein-Protein Interaction Network Construction

To further define potential crosstalk among identified RNA-binding proteins, PPI network was constructed using the STRING database (version 12.0) *(Szklarczyk et al. 2023)*. Both experimentally validated and predicted associations were included, with suitable probabilistic confidence scores applied to every interaction. The STRING web interface was used to determine cluster modules, pathway enrichments, and network topology *(Ono et al. 2025)*. This network-based approach enabled the detection of highly connected RBP clusters and provided a systems-level understanding of lncRNA-protein interactome architecture.

### 2.4 *In Vitro* Transcription

*In vitro* transcription (IVT) was performed using T7 RNA polymerase to generate the PQS of the lncRNA under investigation. The oligonucleotide sequence was carefully designed to accommodate the inherent constraints of T7 polymerase-based transcription *(Singh et al. 2023)*. Specifically, the T7 consensus initiation sequence (GGGAGA) was modified to GCGAGA in the sense strand, as reported in previous studies, to prevent the unintended incorporation of an additional guanine tract and to minimize 5’ heterogeneity. To further mitigate sequence variability, an additional ten-base-pair sequence was incorporated adjacent to the PQS of interest, ensuring minimal disruption to the intended structure, and a twenty-base-pair primer sequence was appended at the 3’ end of the sense strand to facilitate visualization in the reverse transcriptase stop assay. **Figure S1** illustrates the overall design of the DNA templates and the workflow used to generate RNA constructs containing PQS through T7 RNA polymerase mediated *in vitro* transcription.

Oligonucleotides for IVT were procured from Sigma Aldrich Ltd. Bangalore as a dry powder (0.05 mmol scale) and reconstituted in nuclease-free water. Annealing of the IVT template and T7 promotor sequence was in a buffer consisting of 10 mM Tris pH 8.0, 50 mM NaCl, and 1 mM EDTA pH 8.0 at 95 °C for five minutes, followed by a gradual cool-down (∼2 hours) to room temperature. The annealed oligonucleotides were quantified using a NanoDrop™ 2000c Spectrophotometer. Following the manufacturer’s protocol, IVT was conducted using the HiScribe® T7 High Yield RNA Synthesis Kit (New England Biolabs). RNase Inhibitor Murine was used to prevent degradation. Following IVT, RNA purification was performed using the Monarch® RNA Cleanup Kit (New England Biolabs), and the purity and concentration were assessed via NanoDrop™. To prevent degradation, RNAse Inhibitor Murine was added to the purified RNA.

### 2.5 Native PAGE

Native PAGE was performed directly after IVT, by first mixing RNA (2 µM) with loading dye in nuclease-free water (20 µL total). For PAGE with increasing concentration of ions, the RNA (2 µM) was folded in a buffer consisting of 10 mM Tris-Cl (pH 7.5) and 0.1 mM EDTA (pH 8.0) by heating at 95 °C for 5 minutes and then gradually cooled down to room temperature in the presence of increasing concentration of K^+^ or Li^+^. Samples were electrophoresed using 15% polyacrylamide gel at 120 V for ∼1.5 h in 1X TBE using the BioRad Mini-PROTEAN system. Gels were rinsed and imaged using the Bio-Rad ChemiDoc MP system. Staining was performed with Thioflavin T (0.5 mM ThT) for 20 min on a rocker, followed by imaging with the Emerald Pro Q filter (488nm channel). The gel was then stained with Diamond Nucleic Acid Dye for 20 min and imaged using the EtBr filter.

### 2.6 Circular Dichroism Spectroscopy

RNA samples (4 μM) were folded in a buffer containing 10 mM Tris-Cl (pH 7.5) and 0.1 mM EDTA (pH 8.0) by heating at 95 °C for 5 minutes, followed by slow cooling to room temperature. RNA samples were folded in the presence of 100 mM KCl or LiCl or along with varying concentrations of TMPyP4 (5-30 μM) and 5 µg of HSA, as specified for each experiment. Circular Dichroism (CD) spectra were acquired using a JASCO J-815 spectropolarimeter with a 1 mm quartz cuvette under the following settings: 100 nm/min scan speed, 190-320 nm wavelength range, 1 s DIT, 1.00 nm bandwidth, 3 accumulations, at 16 °C. Spectra were smoothed using the Savitzky-Golay method (15-point window), and mean CD intensities (mdeg) were plotted against wavelength (200-320 nm).

### 2.7 Circular Dichroism Thermal Melting Assay

CD melting experiments were performed using a J-810 spectropolarimeter (JASCO Co., Ltd., Japan) equipped with a Peltier temperature controller using a 1 mm quartz cuvette. Samples were prepared in a buffer containing 10 mM Tris-Cl (pH 7.5) and 0.1 mM EDTA (pH 8.0) by heating at 95 °C for 5 minutes, followed by slow cooling to room temperature. The melting profiles were obtained by monitoring the CD signal at 267 nm, corresponding to the maximal positive ellipticity. The temperature was increased from 20 °C to 95 °C at a heating rate of 1 °C/min. The resulting melting curves were fitted to a sigmoidal Boltzmann model to derive the melting temperature (T_m_), defined as the inflexion point of the transition.

### 2.8 Thioflavin T Fluorescence Assay

RNA samples (2 µM) were prepared in folding buffer (10 mM Tris-Cl, pH 7.5; 0.1 mM EDTA, pH 8.0) in a final volume of 150 µL. Where indicated, samples contained 100 mM KCl or LiCl, or HSA. Control reactions lacking RNA and/or ThT were also included. For protein interaction assays, folded RNA (2 µM) was incubated with increasing concentrations of HSA (0-60 µM) in folding buffer at 37 °C for 2 h. ThT was then added to a final concentration of 2 µM. Each reaction (50 µL) was transferred in triplicate to black 96-well plates, and end-point emission was also measured at 488 nm after excitation at 445 nm. using a Cytation 5 reader (Agilent Technologies, USA). Data from two independent experiments (each performed in triplicate) were analyzed in Microsoft Excel and GraphPad Prism. Fluorescence intensities and fold-change values (F/F₀) were plotted, and statistical significance was determined using ordinary one-way ANOVA.

### 2.9 Reverse Transcriptase Stop Assay

A modified version of the reverse transcriptase stop assay originally described by Hagihara et al. was employed *(Hagihara et al. 2010)*. Texas Red-labeled primers (Sigma-Aldrich, USA) were reconstituted in nuclease-free water to a final concentration of 100 μM. Each reaction (10 μL) contained 2 μM RNA, 200 nM labeled primer, 2 mM dNTPs (Promega, Cat. No. U1511), and annealing buffer (10 mM Tris pH 8.0, 50 mM NaCl, 1 mM EDTA pH 8.0), with varying KCl or LiCl concentrations (0-150 mM). The mixture was denatured at 95 °C for 5 min and then allowed to cool to room temperature for primer annealing. Reverse transcription was initiated by adding 4 U/μL reverse transcriptase (Promega, Cat. No. M5301) along with a buffer containing 3 mM MgCl₂, 10 mM Tris-Cl pH 7.5, and 1 mM EDTA pH 8.0, followed by incubation at 37 °C for 1 hour. Reactions were stopped by adding an equal volume of stop buffer (95% formamide, 0.05% bromophenol blue, 20 mM EDTA, 0.05% xylene cyanol). Products were resolved on a 15% denaturing urea-PAGE gel and imaged using a ChemiDoc MP system with a Rhodamine filter. Gels were then stained with Diamond Nucleic Acid Dye (Promega) to visualize primer bands. Band intensities were quantified using ImageJ, and statistical significance was assessed using one-way ANOVA.

### 2.10 Isothermal Titration Calorimetry

Isothermal titration calorimetry (ITC) was performed using a MicroCal ITC200 instrument (Malvern Panalytical) at 25 °C to investigate the interaction between folded lncRNA and HSA. RNA samples (5 μM) were prepared in folding buffer (10 mM Tris-Cl, pH 7.5; 0.1 mM EDTA, pH 8.0), denatured at 95 °C for 5 minutes, and slowly cooled to room temperature over 2 hours. The folded RNA was loaded into the sample cell, and HSA (10-20 μM) prepared in the same buffer was placed in the syringe, which was rotated at 750 rpm. A total of 20 injections (1-2 μL each) were delivered at 150-second intervals to ensure equilibrium, with the initial 0.4 μL injection excluded from data analysis to minimize artifacts from diffusion. The reference power was set to 10 μcal/sec. Data was analyzed using the MicroCal software, applying a single-site binding model to calculate thermodynamic parameters including the dissociation constant (K_d_), enthalpy (ΔH), entropy (ΔS), and binding stoichiometry (n). The binding profile provided insights into the spontaneity and nature of the interaction based on ΔG and whether the process was endothermic or exothermic.

### 2.11 Electrophoretic Mobility Shift Assay

Electrophoretic Mobility Shift Assay (EMSA) was used to evaluate interactions between *in vitro* transcribed lncRNA PQS and HSA. RNA (4 μM) was folded in a buffer consisting of 10 mM Tris pH 8.0, 50 mM NaCl, 1 mM EDTA pH 8.0 in the presence of Texas Red-tagged primer (200 nM). The mixture was heated to 95 °C for 5 minutes and slowly cooled to room temperature to allow proper folding and primer binding. The labeled RNA was then incubated with increasing concentrations of HSA (0-30 μM) in a 10 μL reaction at 37 °C for 2 hours. Binding reactions were resolved on 8% native polyacrylamide gels (1X TBE), electrophoresed at 120 V for 45 minutes using the Bio-Rad Mini-PROTEAN system. Gels were first imaged using the Rhodamine filter on the Bio-Rad ChemiDoc MP system to visualize the Texas Red-labeled RNA. Subsequently, gels were stained with 0.5 mM ThT prepared in methanol and 1X TBE, incubated for 20 minutes, and imaged using the Pro Q Emerald 488 channel to detect G4 signals.

Band intensities were quantified using ImageJ. Statistical significance across HSA concentrations was assessed using one-way ANOVA, with results expressed as mean ± SEM from three independent experiments. Significance thresholds were set at *p < 0.05, **p < 0.01, ***p < 0.001, and ****p < 0.0001.

### 2.12 Preparation of Simulated Body Fluid (SBF)

We followed the methodology described by Kokubo et al. in 2006 to prepare simulated body fluid (SBF) *(Kokubo and Takadama 2006)*. The ion concentrations used to prepare SBF are provided in Table S1. SBFs are widely used *in vitro* systems that mimic physiological conditions to study drug dissolution, biomaterial bioactivity, and toxicological exposure, improving prediction of in vivo behavior *(Gray et al. 2010; Marques et al. 2011)*. It is particularly important for evaluating implant materials through apatite formation, which correlates with bone-bonding ability and can reduce animal testing *(Zhao et al. 2017)*. Additionally, these fluids help assess environmental risks, such as mercury bioaccessibility from contaminated waste *(Gray et al. 2010)*. G4 folding and stability are highly sensitive to physiological conditions such as ionic composition, pH, and molecular crowding, all of which are more accurately replicated in SBFs than in simple buffers. This enables more realistic assessment of G4 formation improving the physiological relevance and translational value of findings.

## 3. Results and Discussion

### 3.1 Putative G-quadruplex-forming sequences in circulating lncRNAs dysregulated in cancer

The Lnc2Cancer (Version 3.0) database was used to identify 1007 circulating lncRNA entries associated with various cancers. After removal of duplicate entries, previously investigated G4-forming lncRNAs, non-lncRNA or partial sequences, and RefSeq entries classified as predicted or provisional, a final set of 283 validated human lncRNAs was retained for G4 prediction analysis (**Figure 1**). After checking for redundant names in GeneCards, lncRNA sequences were extracted in FASTA format and analyzed using QGRS Mapper and G4Hunter tools. PQS sites were identified and categorized into 2G (232 sequences), 3G (36 sequences), and 4G (4 sequences) based on stability. Based on the scores obtained, loop length, and other factors, we further shortlisted four lncRNAs to perform *in vitro* experiments to validate the in silico findings of G4 formation. The workflow was also run in parallel using the portal www.canlncg4.com which does not have a separately curated set of circulating lncRNAs. These four lncRNA sequences included KRTAP5-AS1 (Keratin Associated Protein 5 Antisense 1) containing one 2G sequence, two 3G sequences containing lncRNA named AGAP2-AS1 (ArfGAP with GTPase domain, ankyrin repeat and PH domain 2 antisense 1) and LINC00683 (Long Intergenic Non-Coding RNA 00683) and one 4G sequence containing lncRNA named DLG1-AS1 (Discs Large MAGUK scaffold protein 1 antisense RNA 1). The detailed in silico pipeline for finding these four lncRNAs is illustrated in **Figure 1**. The expression patterns of the selected circulating lncRNAs, their associated cancers and PQS scores are summarized in **Table 1**.

**Figure 1.**
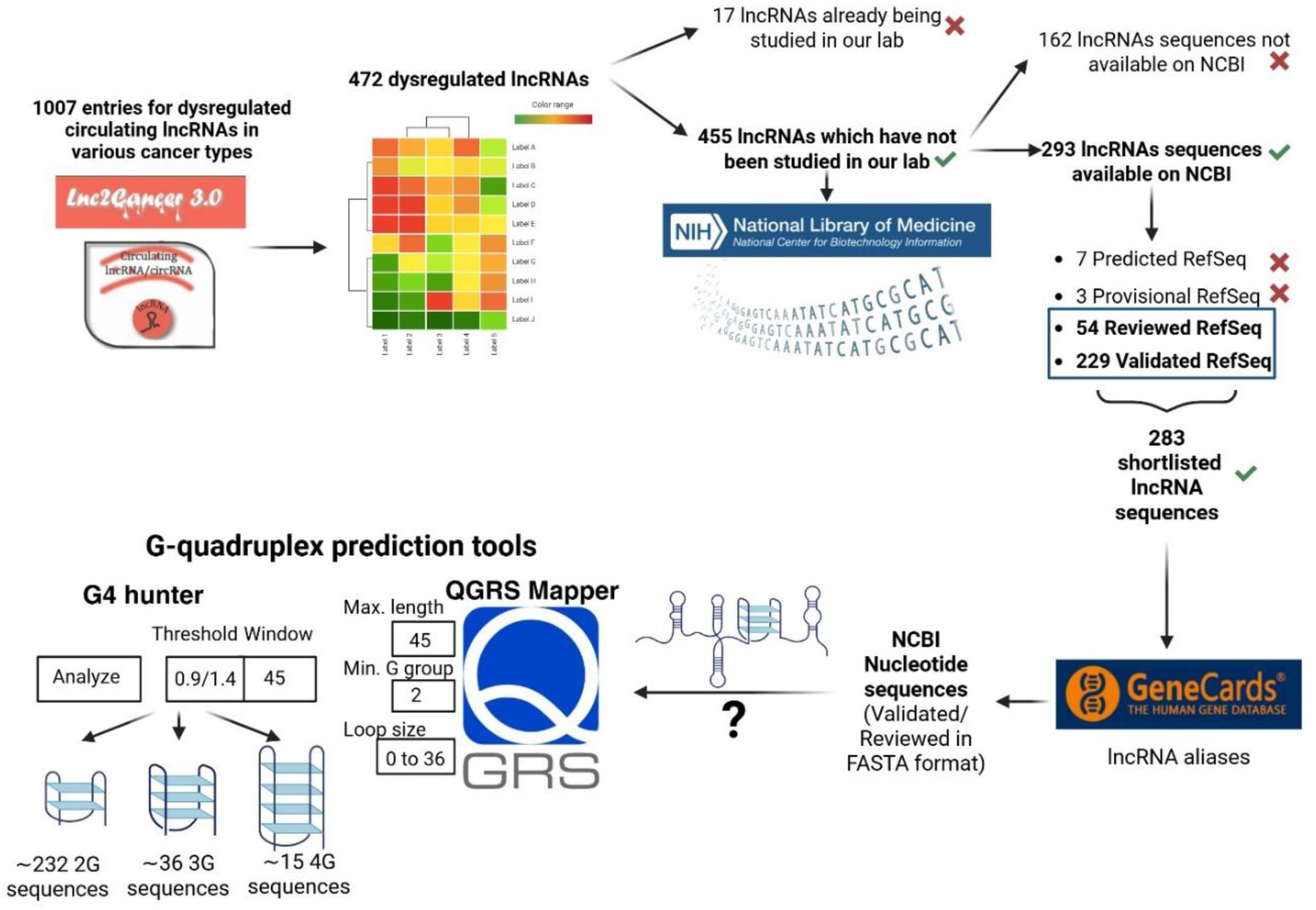
Workflow for computational identification of circulating lncRNAs which harbor G4 motifs.

**Table 1.**
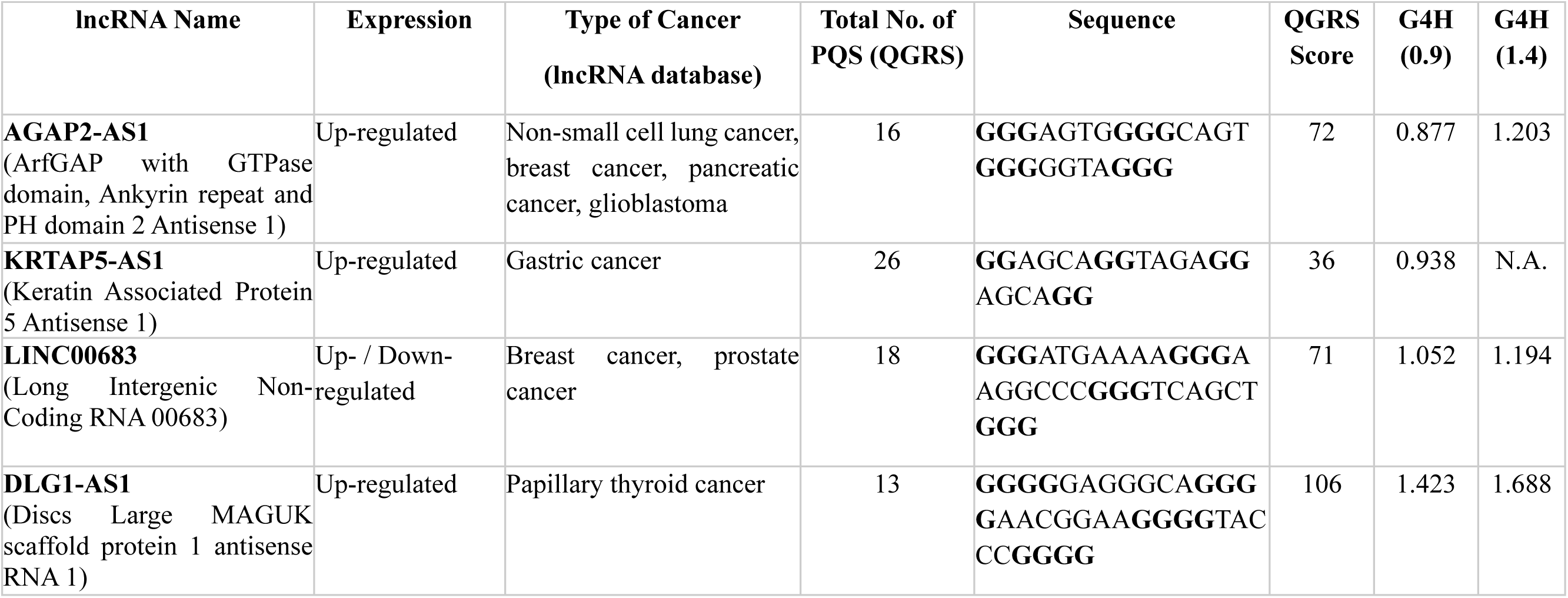
List of selected G4 bearing circulating lncRNAs.

### 3.2 Identification and interaction prediction of G4-Binding RNA-Binding Proteins (RBPs) associated with circulating lncRNAs

The LncTarD and NPInter databases were used to retrieve the experimentally supported and predicted protein interaction profiles of the circulating lncRNAs. LncTarD database also yielded a network analysis **(Figure S2)** demonstrating AGAP2-AS1 as a central regulatory hub in cancer. Previous reports have shown AGAP2-AS1 as being transcriptionally activated by SP1 transcription factor in gastric cancer, and ineracting with proteins such as ELAVL1 (HuR), EZH2, CDH1, NOTCH2, MMP2, and ANKRD1 in breast, lung, and gastrointestinal cancers *(Ji et al. 2021)(Luo et al. 2019)*. AGAP2-AS1 also has a validated role as a competitive endogenous RNA (ceRNA) sponging various miRNAs including miR-16-5p, miR-424-5p, miR-182-5p, miR-15a/b-5p, and miR-296, across different cancers *(Lu et al. 2024)*.

We next sought to isolate RBPs that possibly mediate G4 structure recognition in the selected lncRNAs. Towards this objective, the protein lists from both the databases for the selected lncRNAs (AGAP2-AS1, DLG1-AS1, KRTAP5-AS1, LINC00683) were checked against a list of 24 proteins with validated G4-binding activity from the G4IPDB database. This integrative analysis revealed ten proteins; ELAVL1, SP1, DHX36, IGF2BP1, hnRNPA2B1, FUS, HNRPK, ILF3, SFRS1, and FMR1; to interact with one or more of these lncRNAs. These RBPs are of particular interest given their relevance in RNA structure regulation, including G4 motif binding, and provide a rational set of candidates for further mechanistic validation (**Table 2**).

**Table 2.** List of RNA-binding proteins with G4 interactions for lncRNAs.

| Protein Name | lncRNA | Reference |
| --- | --- | --- |
| ELAVL1 | AGAP2-AS1 | (*Han et al. 2022*) |
| SP1 | AGAP2-AS1, DLG1-AS1, KRTAP5-AS1, LINC00683 | (*Qi et al. 2017*) |
| DHX36 | AGAP2-AS1, KRTAP5-AS1, LINC00683 | (*Sauer et al. 2019*) |
| IGF2BP1 | AGAP2-AS1, DLG1-AS1, LINC00683 | (*Elcheva et al. 2020*) |
| hnRNPA2B1 | DLG1-AS1, KRTAP5-AS1, LINC00683 | (*Liu et al. 2022*) |
| FUS | DLG1-AS1, KRTAP5-AS1, LINC00683 | (*Lagier-Tourenne et al. 2012*) |
| HNRPK | KRTAP5-AS1 | (*2012*) |
| ILF3 | KRTAP5-AS1 | (*2012*) |
| SFRS1 | KRTAP5-AS1 | (*2012*) |
| FMR1 | KRTAP5-AS1 |  |

Footnote: All interactions were retrieved from the NPInter database unless otherwise specified. ELAVL1 interaction data was obtained from lncTarD, while SP1 interactions were supported by both lncTarD and NPInter.

To evaluate the direct interaction propensities between these G4-binding RBPs and the selected lncRNAs, we performed in silico RPISeq predictions using both RF and SVM classifiers. Only protein-lncRNA pairs with scores exceeding the 0.5 confidence threshold by both methods were considered strong potential interactors. This two-tiered workflow integrates curated experimental data and computational assessment, ensuring that downstream analyses focus specifically on high-probability, functionally significant lncRNA-protein interactions that are enriched for G4 regulatory activity. The RPI-Seq interaction scores for circulating lncRNAs and G4-binding RBPs is shown in **Table S2**. DHX36, a well-characterised G4-resolving RNA helicase, showed high interaction probabilities with AGAP2-AS1, KRTAP5-AS1, and LINC00683, supporting its role as a shared interactor among G4-containing lncRNAs involved in post-transcriptional regulation *(Tippana et al. 2019)*. In contrast, ALB, FUS, IGF2BP1, hnRNPA2B1, and SP1 displayed differential interaction patterns between LINC00683 and DLG1-AS1, reflecting differences in their structural and functional properties. LINC00683 preferentially interacted with IGF2BP1, hnRNPA2B1, and FUS, consistent with its stable GC-rich structured architecture and putative m6A sites, whereas DLG1-AS1 showed higher affinity for SP1 and hnRNPA2B1, aligning with its partially unstructured antisense organization *(Thompson et al. 2023; Alarcón et al. 2015; Duan et al. 2024a; Nguyen et al. 2018; Liu et al. 2022; Huang et al. 2018; Liu and Xiang 2023)*. Collectively, these findings suggest that while DHX36 functions as a common structure-specific helicase across multiple G4-containing lncRNAs, other RBPs contribute more selectively to RNA stabilisation, transcriptional regulation, and higher-order ribonucleoprotein complex formation *(Richardson et al. 2020)*.

### 3.3 Protein-Protein Interaction Network based on PQS-containing lncRNAs

Since our shortlisted circulating lncRNAs contain PQS and are enriched for G4-binding RBPs, we constructed a high-confidence STRING-based (confidence threshold: 0.7) PPI network to identify cooperative protein modules and functional hubs underlying lncRNA stability, RNA metabolism, and cancer progression.

The network in **Figure S3** includes 11 nodes; ELAVL1, IGF2BP1, hnRNPA2B1, FUS, hnRNPK, ILF3, SRSF1, FMR1, SP1, ALB, and DHX36; and 22 edges, with an average node degree of 4 and a strong average local clustering coefficient of 0.633. Statistical evaluation returned a PPI enrichment p-value less than 1.0e-16, indicating that these proteins interact far more frequently than would be expected by chance, reflecting their shared biological roles in post-transcriptional gene regulation *(Szklarczyk et al. 2023)*. To further resolve the modular structure within this network, we applied the Markov Cluster Algorithm (MCL); a graph-based approach optimized for biological network data *(Moschopoulos et al. 2011)*. This analysis identified a single dominant cluster encompassing eight core RBPs (hnRNPA2B1, IGF2BP1, ELAVL1, FMR1, hnRNPK, FUS, ILF3, SRSF1). The remaining proteins (ALB, DHX36, SP1) remained outside this core, consistent with more peripheral or context-specific roles.

The identified cluster was strongly enriched for “negative regulation of mRNA metabolism,” mirroring the results of our Gene Ontology (GO) enrichment analysis. GO enrichment analysis (**Figure 2**) for this RBP network reveals a consistent and statistically robust overrepresentation of RNA metabolic processes. The most significant biological processes include negative regulation (FDR = 9.19e-08) and general regulation (FDR = 5.33e-07) of mRNA metabolic process, negative regulation of RNA catabolic process (FDR = 2.95e-06), RNA stabilisation (FDR = 7.76e-05), and negative regulation of mRNA catabolic process (FDR = 7.85e-05). These results indicate a central role in controlling mRNA decay, stability, and translation. Enrichment for molecular functions such as mRNA binding (FDR = 2.80e-11), mRNA 3’-UTR binding (FDR = 1.04e-10), AU-rich region binding (FDR = 0.00046), mRNA 5’-UTR binding (FDR = 0.00046), and miRNA binding (FDR = 0.00064) indicates direct engagement with mRNA substrates, including regulatory regions mediating transcript decay or stabilisation. Cellular component annotations (e.g., stress granules, ribonucleoprotein complexes, spliceosomal units) reinforce the involvement of these components in RNA granule dynamics and nuclear-cytoplasmic shuttling.

**Figure 2.**
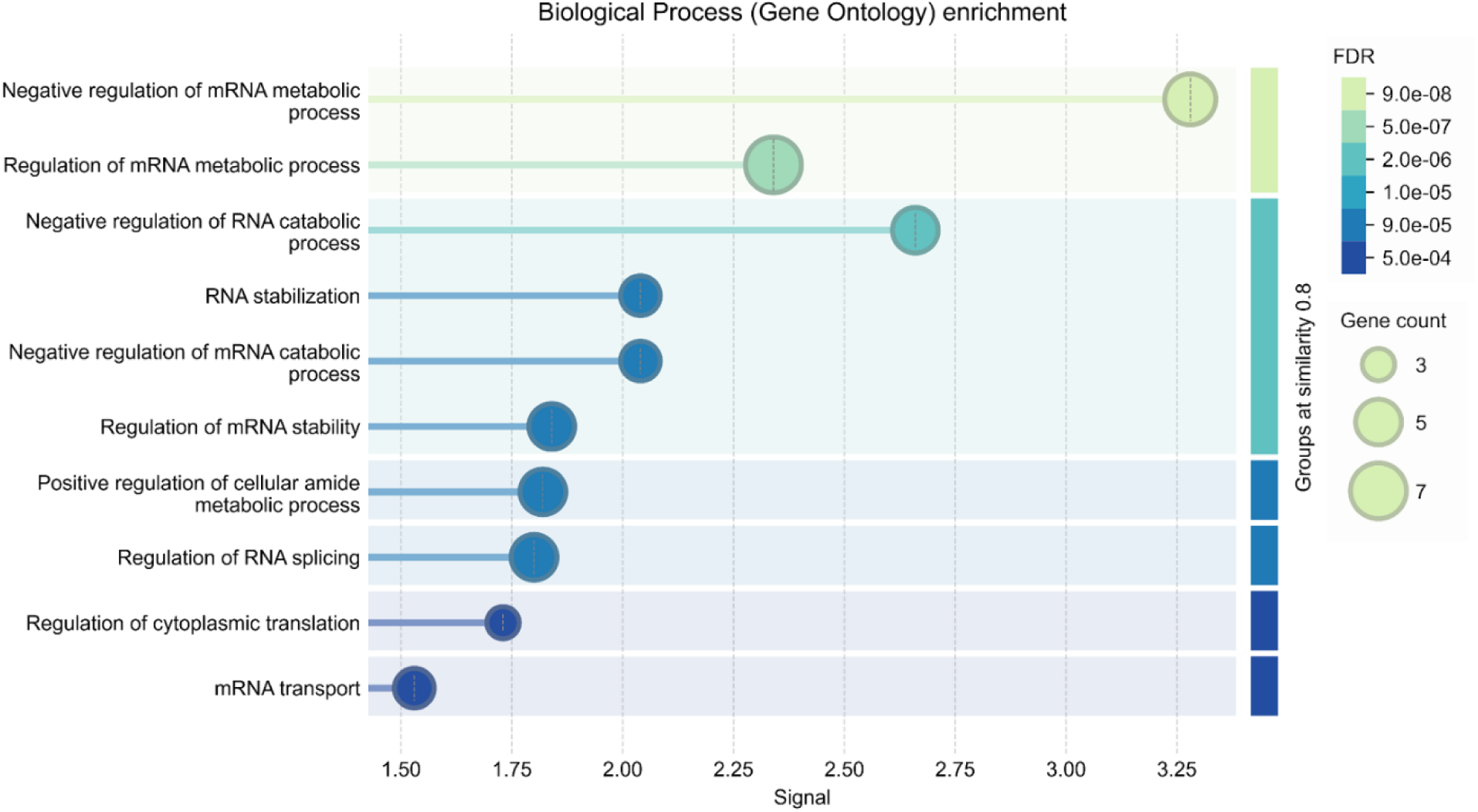
Gene ontology enrichment of biological processes for lncRNA-associated RBPs.

Building on these in silico insights into RNA-binding protein associations and functional enrichment, we next sought to experimentally validate the structural features of the selected lncRNAs, particularly their ability to form G4 conformations and interact with certain proteins *in vitro*.

### 3.4 G4 Formation by PQS of Circulating lncRNAs

Native PAGE analysis performed immediately after *in vitro* transcription, without subjecting the RNA to a folding buffer, revealed two distinct bands for AGAP2-AS1, LINC00683 and DLG1-AS1. This result suggests the presence of multiple RNA conformations, including unfolded or partially folded structures and a small fraction of heterogeneous RNA formed during *in vitro* transcription *(PLEISS et al. 1998; Singh et al. 2025)*. In contrast, upon folding in a buffer consisting 10 mM Tris-HCl (pH 7.5), 0.1 mM EDTA (pH 8.0) via heat denaturation and gradual cooling, a single dominant band was observed with the second band markedly reduced **Figure S4**, suggesting adoption of a more uniform and stable structure, likely a compact G4. This shift reflects a transition from conformational heterogeneity to a single predominant species. In contrast, KRTAP5-AS1 showed no change upon folding, indicating a pre-formed or stable structure. The folding buffer was constituted with 10 mM Tris-HCl (pH 7.5) and 0.1 mM EDTA (pH 8.0).

We investigated the electrophoretic behavior of the selected RNA in presence of K^+^ and Li^+^. As shown in **Figure 3**, with increasing K^+^ concentrations a shift was observed from monomeric to slower-migrating multimeric G4 species. Interestingly, analogous observations were also recorded in presence of Li^+^. While K^+^ ions are generally more effective stabilizers of G4s due to their optimal ionic radius, the precise relationship between cation type, concentration, and the resulting G4 topology is complex and sequence-dependent *(Chen 1992)(Saintomé et al. 2016)*. AGAP2-AS1, DLG1-AS1 and LINC00683 displayed higher amounts of slower-migrating species at lower concentrations of K^+^ and Li^+^, that are consistent with intermolecular G4 assembly. Interestingly, KRTAP5-AS1 displayed no cation-dependent change in molecularity, likely reflecting a structurally rigid or predominantly intramolecular G4 conformation. The short length and limited G-tract content of the KRTAP5-AS1 PQS may further restrict multimeric G4 formation.

**Figure 3.**
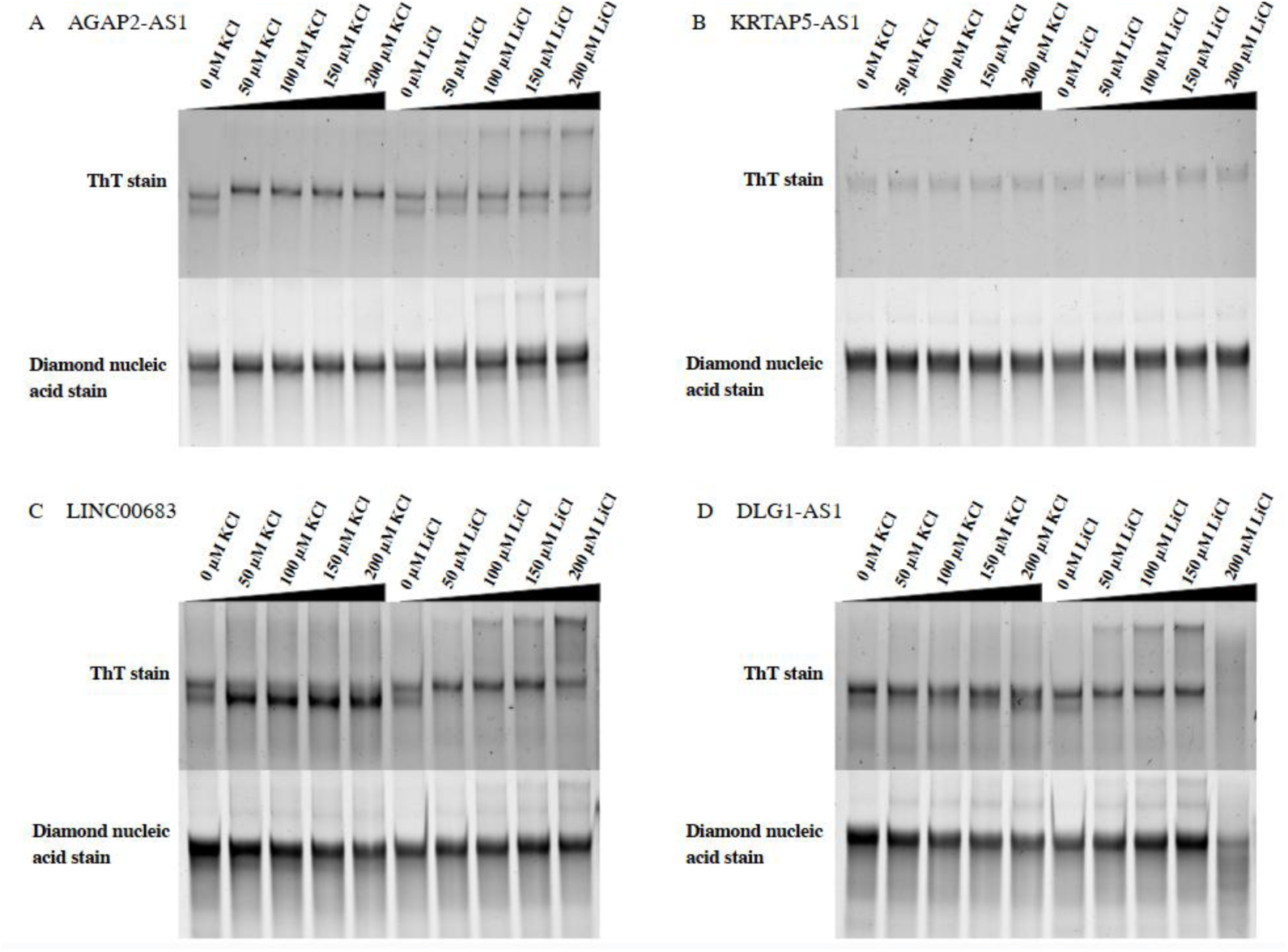
Native polyacrylamide gel electrophoresis of (A) AGAP2-AS1, (B) KRTAP5-AS1, (C) LINC00683, and (D) DLG1-AS1 under increasing concentrations of KCl and LiCl. Gels were visualized using ThT staining to detect G4 structures, followed by counterstaining with diamond nucleic acid dye for total RNA visualization.

The differential effects of K^+^ and Li^+^ highlight the nuanced role of cation size and charge density in dictating the equilibrium between monomeric and multimeric G4 architectures, a key aspect of G4 structural plasticity. CD spectroscopy demonstrated that all four IVT-derived lncRNAs form parallel G4 structures, evidenced by a positive peak around ∼265 nm and a negative peak near ∼240 nm, along with an additional characteristic positive peak at ∼210 nm, observed both in the absence and presence of monovalent cations (100 mM KCl or LiCl) (**Figure 4 A-D**). These trends are consistent with EMSA observations. CD thermal melting analysis **(Figure 4 E)** showed that AGAP2-AS1 displayed a clear transformation from folded to unfolded state. In contrast, the CD thermal melting of KRTAP5-AS1, LINC00683 and DLG1-AS1 revealed incomplete transition to unfolded state.

**Figure 4.**
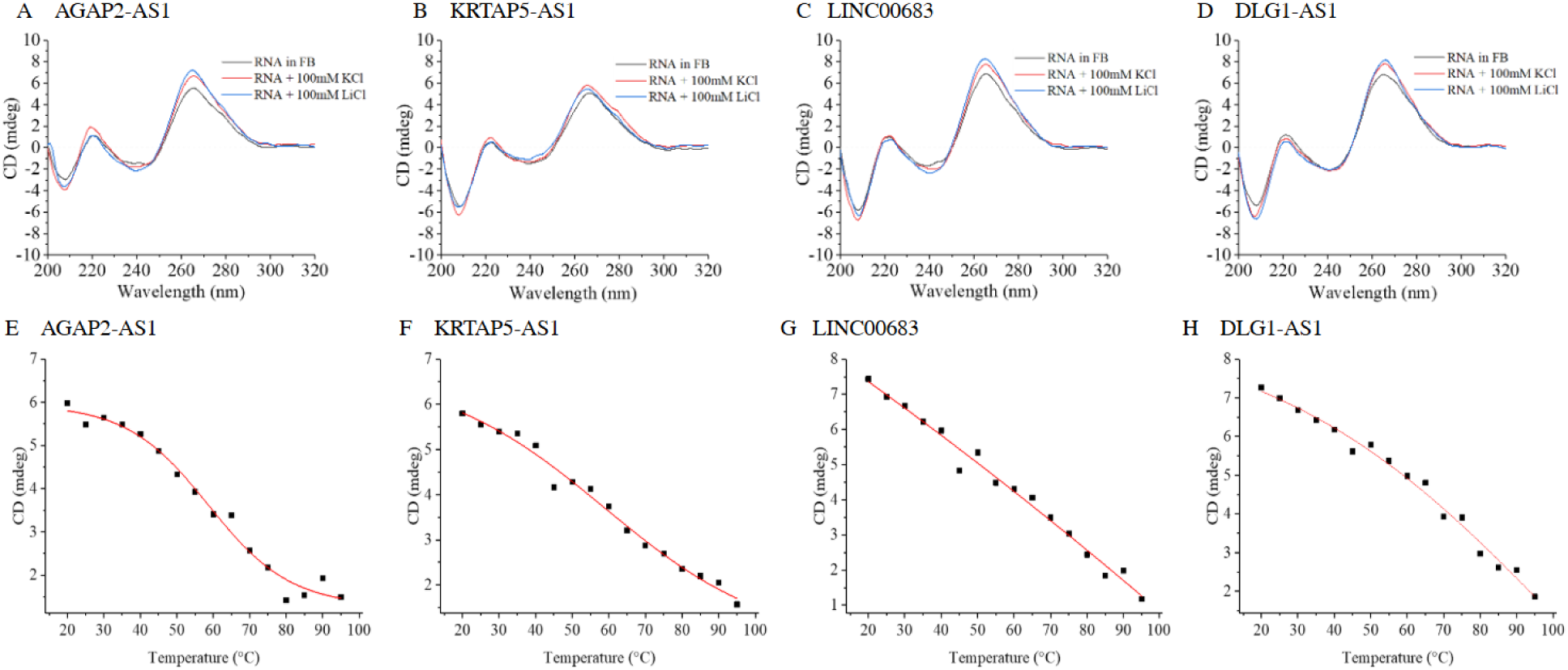
Circular dichroism spectra and thermal melting of PQS. (A-D) CD spectra of PQS (4 μM) A. AGAP2-AS1, B. KRTAP5-AS1, C. LINC00683, D. DLG1-AS1 recorded in folding buffer alone and in the presence of monovalent cations K⁺ (100 mM) or Li⁺ (100 mM) to assess ion-dependent G4 folding. (E-H) CD melting curves demonstrating changes in the thermal stability of E. AGAP2-AS1, F. KRTAP5-AS1, G. LINC00683, H. DLG1-AS1 recorded in folding buffer alone.

We next studied the effect of TMPyP4 on the PQS. TMPyP4 is a porphyrin ligand that is known to destabilize RNA G4s (**Figure 5 A-D**) (*Morris et al. 2012; Zamiri et al. 2014)*. CD spectra of all the PQS in presence of increasing amounts of TMPyP4 show some loss of the CD maxima and minima at 265 nm and 240 nm, respectively. The reduction in CD signal intensity is not uniform across the RNAs with a noticeably greater disruption observed for AGAP2-AS1 compared to only modest loss for KRTAP5-AS1 and LINC00683. While the loss of CD band intensity for the RNAs is most pronounced at higher concentrations of TMPyP4, DLG1-AS1 appears to be the least affected in presence of the ligand. Overall, these results indicate the persistence of the RNA G4s under destabilizing conditions. We performed EMSA on the RNA PQS in presence of TMPyP4 to check for ligand effects in a different experimental format. EMSA results reflected the relatively stronger G4 motif of DLG1-AS1 compared to KRTAP5-AS1, AGAP2-AS1, and LINC00683 (**Figure 5 E-H**). While the EMSA results are consistent with the findings of CD spectroscopy, the superior sensitivity of the former is evident from the observation of effects at a lower concentration range of TMPyP4.

**Figure 5.**
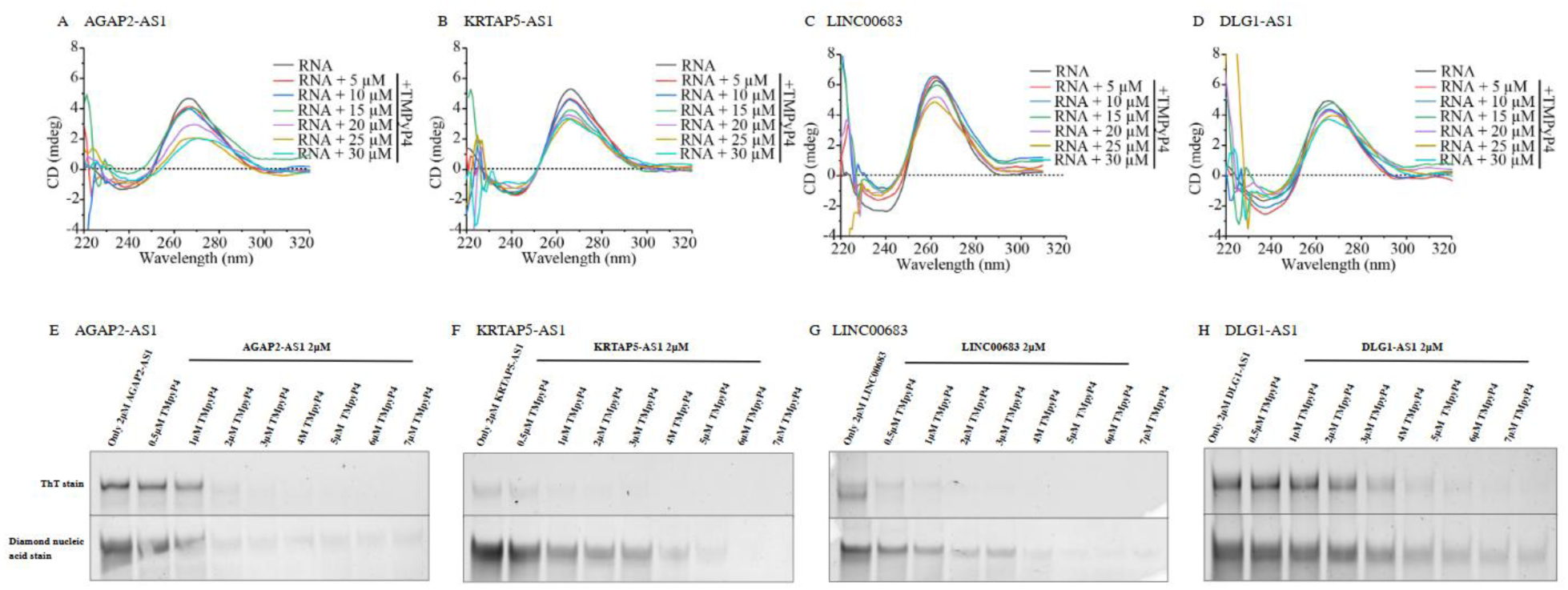
Effect of TMPyP4 on G4 structure and mobility of PQS. (A-D) CD spectra of PQS. (A) AGAP2-AS1, (B) KRTAP5-AS1, (C) LINC00683, and (D) DLG1-AS1 recorded in the presence of increasing concentrations of the G4 ligand TMPyP4 (5, 10, 15, 20, and 25 μM); (E-H) EMSA of (E) AGAP2-AS1, (F) KRTAP5-AS1, (G) LINC00683, and (H) DLG1-AS1 incubated with TMPyP4 at concentrations ranging from 0.25 to 7 µM.

CD spectroscopic or gel electrophoretic methods are limited by their inability to capture the dynamic character of G4s. To this end, we used the reverse transcriptase (RT) stop assay to assess the behavior of the select PQS towards an actively processing enzyme. This polymerase stalling technique relies on the ability of RT to pause or terminate at G4s resulting in truncated cDNA products that can be detected by denaturing PAGE. In the RT stop assay conducted for AGAP2-AS1, DLG1-AS1, and LINC00683, heterogeneous template bands and multiple RT stop sites were observed, resulting in smeared or complex banding patterns that were challenging to interpret. This is likely due to the ability of these RNAs to adopt diverse G4 conformations ranging from monomeric to dimeric or higher-order structures depending on the cationic environment. Such structural heterogeneity can cause RT to stall at various points along the RNA, complicating the identification of specific G4-related stops (**Figure 6**). In contrast, the RT stop assay for KRTAP5-AS1 (**Figure 6B**) did not show any significant change in stop product formation with increasing K⁺ concentrations, suggesting that K⁺ does not markedly influence the stability of the G4 structure formed by this RNA. Interestingly, the addition of Li⁺ led to a noticeable increase in stop products, indicating a stabilizing effect under these conditions (quantified bands in **Figure S5).** This unusual insensitivity toward K⁺ may reflect the inherently stable nature of the G4 formed by KRTAP5-AS1, which appears to maintain its folded state regardless of typical cationic variations. Although rare, some studies have reported deviations from the typical stability order (K⁺ > Na⁺, NH₄⁺, Rb⁺ ≫ Li⁺, Cs⁺). The observed behavior of KRTAP5-AS1 in the presence of Li⁺ likely points to the formation of an atypical G4 structure with an unusual ion-binding preference *(Saintomé et al. 2016; Singh et al. 2025)*.

**Figure 6.**
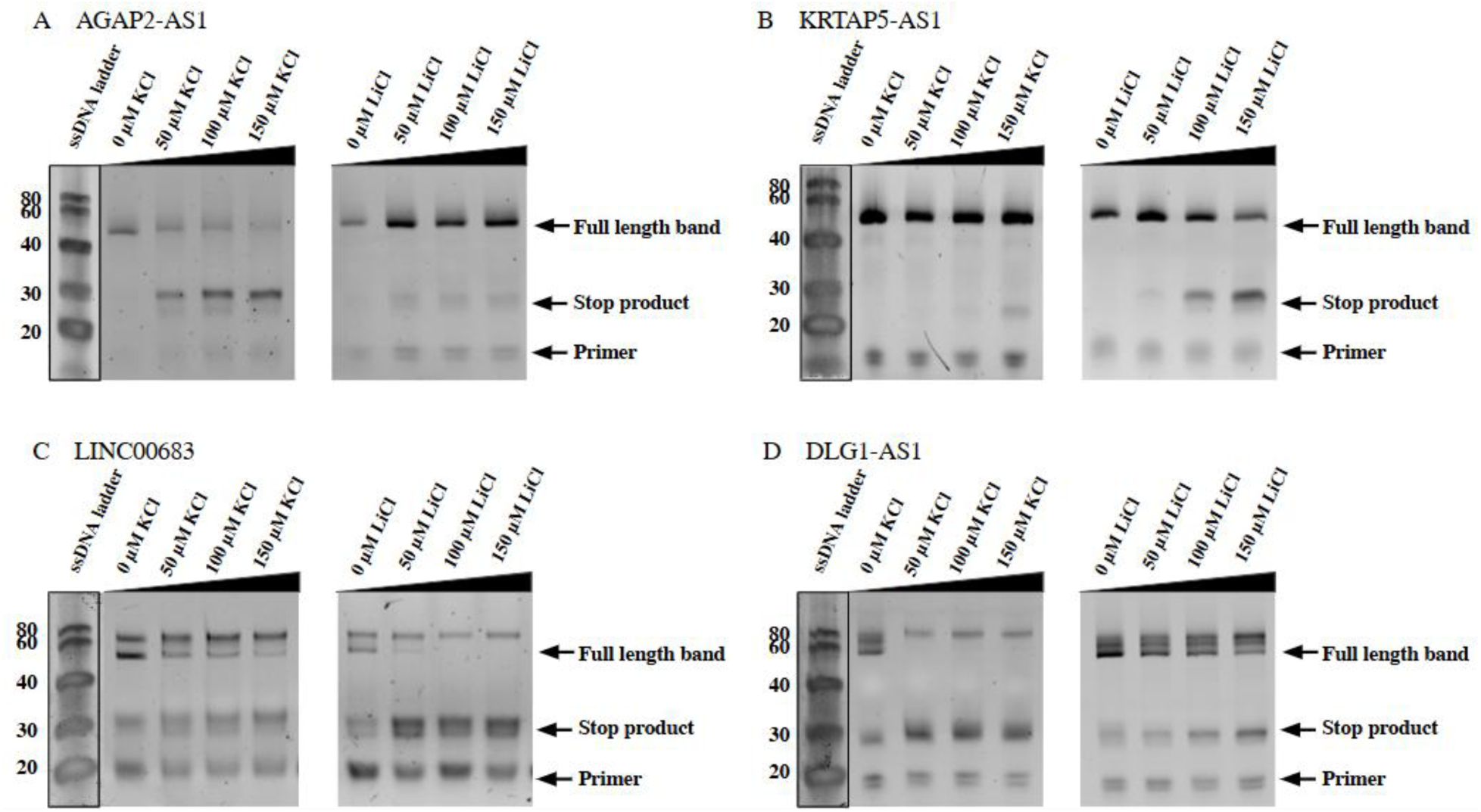
RT-stop assays of (A) AGAP2-AS1, (B) KRTAP5-AS1, (C) LINC00683, and (D) DLG1-AS1 under increasing concentrations of KCl and LiCl. cDNA products were resolved on denaturing PAGE alongside a ssDNA ladder. Full-length bands, stop products, and primer bands are indicated.

### 3.5 EMSA and ITC for HSA binding by PQS

We performed EMSA to examine the interactions between HSA and the PQS. For AGAP2-AS1 (4 μM), increasing HSA concentrations resulted in loss of the free RNA band and appearance of a faster-migrating shifted band, which saturated at ∼15 µM HSA. KRTAP5-AS1, LINC00683, and DLG1-AS1 showed similar behavior, with complex formation reaching saturation at ∼10-15 µM HSA. The faster migration of the RNA-HSA interactions likely arise from HSA-induced conformational changes in the cognate G4s. ThT staining was used to assess G4 integrity within these complexes (**Figure 7**). KRTAP5-AS1 and LINC00683 showed no detectable ThT-stained bands, suggesting a disruption of their G4 structures upon HSA interaction. While AGAP2-AS1 and DLG1-AS1 retained ThT-stained bands in presence of HSA, the intensity of the latter was reduced, suggesting partial destabilization. Based on the EMSA of the RNA PQS in presence of HSA it can be inferred that AGAP2-AS1 and DLG1-AS1 preserve their G4 architecture more effectively than KRTAP5-AS1 and LINC00683. These varied outcomes likely reflect differences in intrinsic G4 stability and topology, pointing to structure-specific modes of RNA-protein interaction.

**Figure 7.**
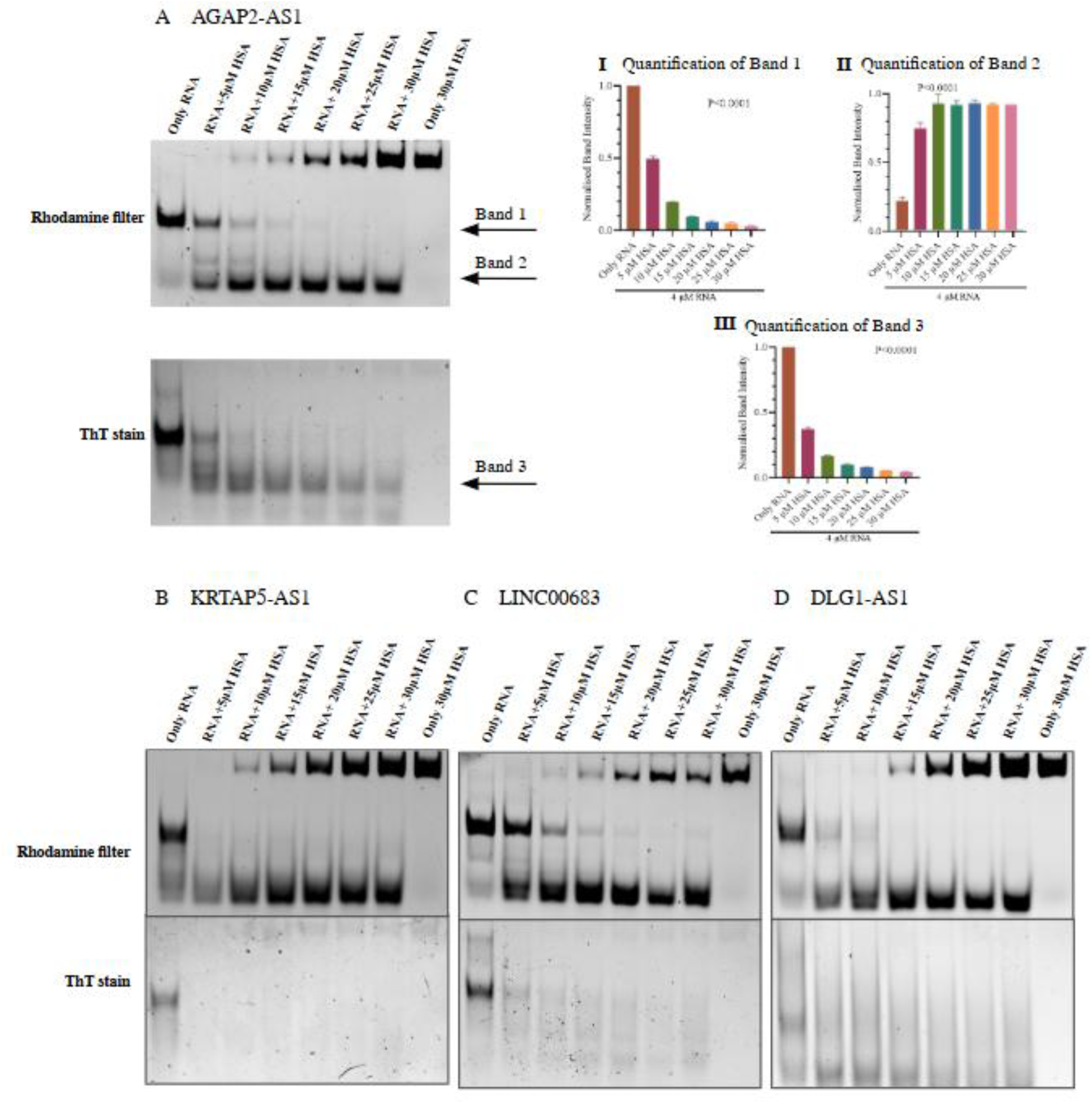
EMSA of HSA interactions with G4-forming lncRNAs (A) EMSA of AGAP2-AS1 (4 μM) with increasing concentration of HSA visualized in rhodamine filter. I, II, III, Quantification of bands 1, 2, and 3, respectively, as indicated in the figure. EMSA of lncRNA B. KRTAP5-AS1, C. LINC00683, and D. DLG1-AS1 with HSA visualized in rhodamine filter followed by ThT staining. Ordinary one-way ANOVA was employed for statistical analysis, and all the resulting P-values were significant.

**Figure 8.**
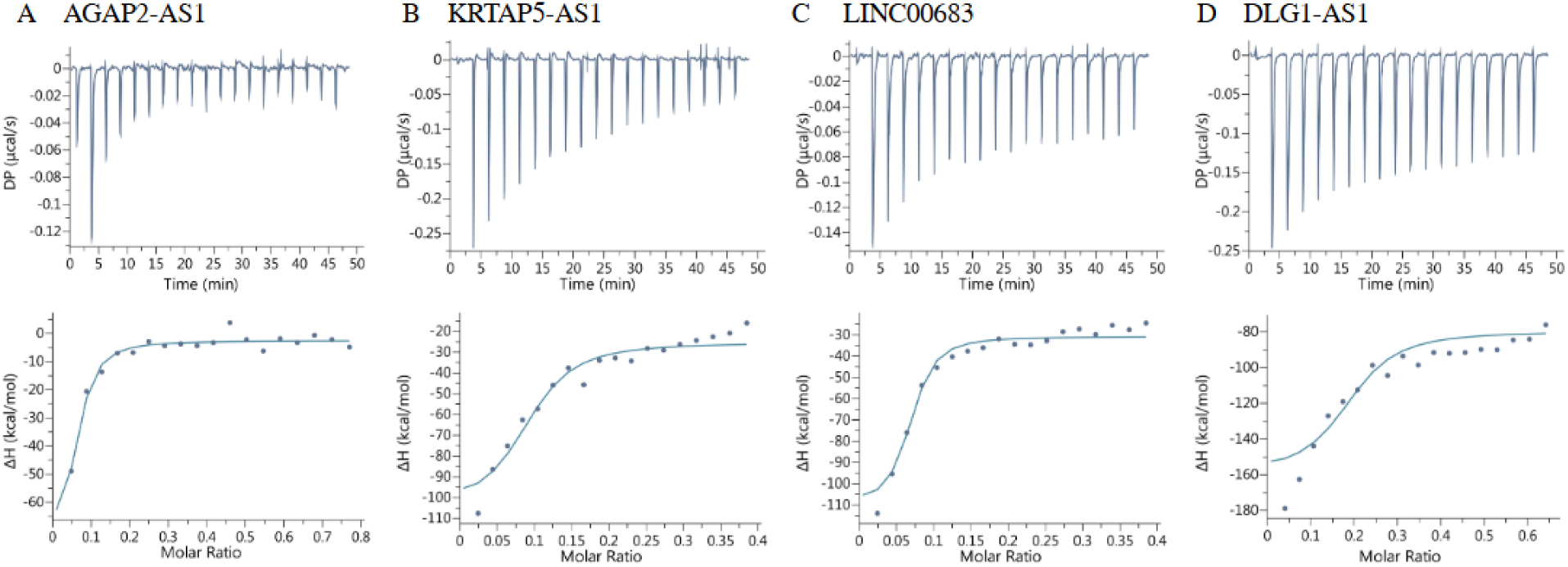
Binding isotherms of A. AGAP2-AS1, B. KRTAP5-AS1, C. LINC00683, D. DLG1-AS1 for the interaction of the RNAs with HSA.

To verify whether the PQS undergo self-association independent of protein interactions, native PAGE was performed across a tenfold RNA concentration range (0.25-5 µM). Gels were sequentially stained with ThT to detect G4 structures and diamond stain to visualize total RNA. For all PQS, a single discrete band was observed at all concentrations, with increased intensity corresponding only to higher RNA loading **(Figure S6)**. No additional shifted bands or smearing were detected, indicating that these lncRNAs form monomeric G4 structures that remain structurally stable and do not self-associate into higher-order complexes under the conditions tested.

We investigated the interaction of the PQS with HSA via ITC. These ITC profiles do not show clear biphasic behavior. Each binding isotherm appears predominantly single sigmoidal, consistent with a one-site or single dominant binding mode rather than two distinct binding events. The four PQS’ displayed comparable binding affinities (K_d_) towards HSA within the same order of magnitude (**Table S3)**. The remarkable nanomolar affinity of the PQS’ towards HSA is distinctive considering the typical K_d_ reported for RNA-protein interactions *(Hudson et al. 2014; Dey et al. 2023; Feig 2009; Haldar et al. 2022)*. Interestingly, LINC00683 displays the strongest K_d_ with HSA (19.4 nM) and yet ThT staining post-EMSA suggests that HSA either unfolds the corresponding G4 or masks it from interaction with ThT. The tightest interaction exercised by LINC00683 with HSA may be accompanied by unfolding of the resident G4 motif that also results in exposure of extra contacts. In contrast, AGAP2-AS1 and DLG1-AS1 bind HSA but the interaction is not accompanied by loss of G4 structure. Notably, KRTAP5-AS1 displays an intermediate K_d_ but greater loss of ThT fluorescence post-EMSA.

High-affinity binding to HSA in the low-nanomolar K_d_ range has been previously reported for dendritic alkyl chain-assisted DNA amphiphiles *(Lacroix et al. 2017)*. However, the high-affinity was driven by hydrophobic modification rather than by a nucleic acid secondary structure per se. The exothermic interactions of all the lncRNA PQS’ with HSA are enthalpy driven and accompanied by a large entropic penalty suggesting significant structural ordering upon complex formation. This thermodynamic feature strengthens the case for a specific, structure-mediated RNA-protein interaction, as opposed to the non-specific electrostatic contacts typical for RNA-polyelectrolyte interactions with albumin reported in earlier literature.

The ThT fluorescence enhancement assay was employed to evaluate the formation, stability, and conformational modulation of G4 structures in IVT-derived PQS under varying monovalent cation conditions and during interaction with HSA. ThT (2 µM) was added post-folding, and end-point emission was recorded at 488 nm after excitation at 445 nm. Relative to RNA-only controls (**Figure 9A**), all PQS exhibited substantial fluorescence enhancement, confirming the presence of G4 structures. Among them, AGAP2-AS1, DLG1-AS1, and LINC00683 showed the highest enhancement (∼35-fold), whereas KRTAP5-AS1 displayed a comparatively lower increase (∼15-fold), suggesting differences in G4 topology or ThT accessibility.

**Figure 9.**
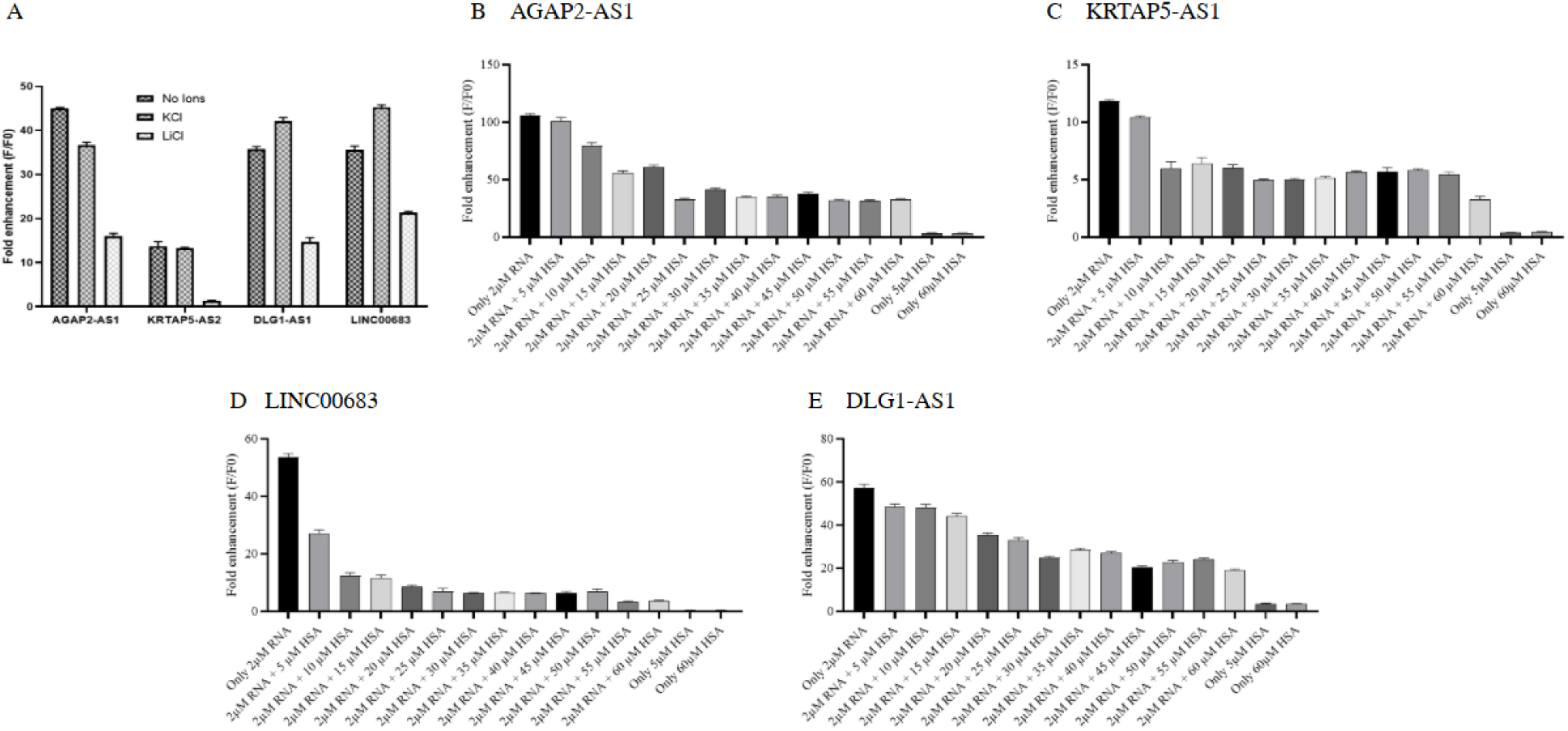
ThT fluorescence enhancement assay showing the effects of monovalent cations and HSA on PQS. A. Fold enhancement of AGAP2-AS1, KRTAP5-AS1, LINC00683, and DLG1-AS1 in the presence/absence of monovalent cations. Fluorescence enhancement (F/F0) plotted against the increasing concentration of HSA and constant concentration of B. AGAP2-AS1, C. KRTAP5-AS1, D. LINC00683, E. DLG1-AS1. Ordinary one-way ANOVA was employed for statistical analysis, and all the resulting P-values were significant.

In the presence of K⁺, LINC00683 and DLG1-AS1 demonstrated increased fluorescence relative to the no-ion condition, consistent with K⁺-mediated stabilization of their G4 structures. In contrast, AGAP2-AS1 and KRTAP5-AS1 exhibited reduced fluorescence enhancement upon K⁺ addition, indicating cation-induced conformational rearrangements that may restrict ThT binding despite G4 stabilization. Under Li⁺ conditions, all PQS showed a further decrease in fluorescence compared to both no-ion and K⁺ environments.

To further assess the responsiveness of PQS to protein interaction, ThT fluorescence assays were performed in the presence of increasing concentrations of HSA (**Figure 9 B-E**). All four lncRNAs, KRTAP5-AS1, AGAP2-AS1, LINC00683, and DLG1-AS1, exhibited a decrease in ThT fluorescence upon HSA addition. This inverse relationship suggests that HSA either competes with ThT for G4 binding due to higher affinity or induces structural rearrangements within the G4 motifs that hinder ThT association. In presence of increasing amounts of HSA, AGAP2-AS1 and DLG1-AS1 display a relatively steady decline in ThT fluorescence (**Figure 9B and E**). In contrast, KRTAP5-AS1 exhibits a plateauing of the ThT fluorescence suggesting the possibility that a portion of the RNA’s structure remains resistant to HSA displacement across a wide concentration range. LINC00683 shows a pronounced decrease in ThT fluorescence enhancement with increasing amounts of HSA suggesting a highly sensitive G4 motif that experiences disruption upon interaction with the protein. These results demonstrate that monovalent cations and HSA modulate G4 conformations in a sequence-dependent manner underscoring the dynamic structural plasticity of the G4s in response to different environmental conditions.

### 3.6 Investigating the Folding of lncRNA PQS in Simulated Body Fluid

Simulated body fluid (SBF) mimics blood plasma in terms of ionic composition (Na⁺, K⁺, Ca²⁺, Mg²⁺, Cl⁻, HCO₃⁻, SO₄²⁻, etc.) and physiological pH (∼7.4), providing a controlled and reproducible *in vitro* system to approximate extracellular and circulating conditions. To evaluate G4 stability in the selected PQS, we employed ThT fluorescence enhancement assays under standard folding buffer and SBF conditions, with and without HSA. The ThT fluorescence enhancement for each of the PQS in absence of HSA is comparable between folding buffer and SBF. Notably, the fold of ThT fluorescence enhancement decreases in the presence of HSA in both folding buffer and SBF albeit to a lesser extent in the latter. The main inference that can be drawn from these results is that the complex ionic environment of SBF confers greater stability to the RNA G4s compared to folding buffer, which in turn provides greater resistance towards HSA-induced conformational changes in the nucleic acid secondary structures. In other words, SBF provides a stabilizing environment that preserves RNA G4 architecture despite the presence of proteins.

Next, we performed CD spectroscopic measurements on the PQS in presence of HSA under folding buffer/SBF conditions. The resulting CD spectra are presented in **Figure S7**. The CD spectra of HSA alone in SBF and folding buffer display intense negative peaks at 208 nm and 220 nm, which are characteristic of an α-helix in the protein. The CD of PQS with HSA in folding buffer/SBF, is dominated by the pronounced CD bands of HSA especially in the wavelength range of 210 – 220 nm. Thus, the CD spectrum of native HSA was subtracted from the total RNA + HSA spectra to obtain corrected RNA-specific curves and enable clearer interpretation of lncRNA conformation. Comparison of the HSA-subtracted CD spectra in folding buffer and SBF (**Figure 10E – L**) clearly indicate the disruption of G4 motifs in each lncRNA. Interestingly, the extent of structural perturbation in the G4s due to interaction with HSA is comparable between folding buffer and SBF **(Figure S8 and Figure S9)**.

**Figure 10.**
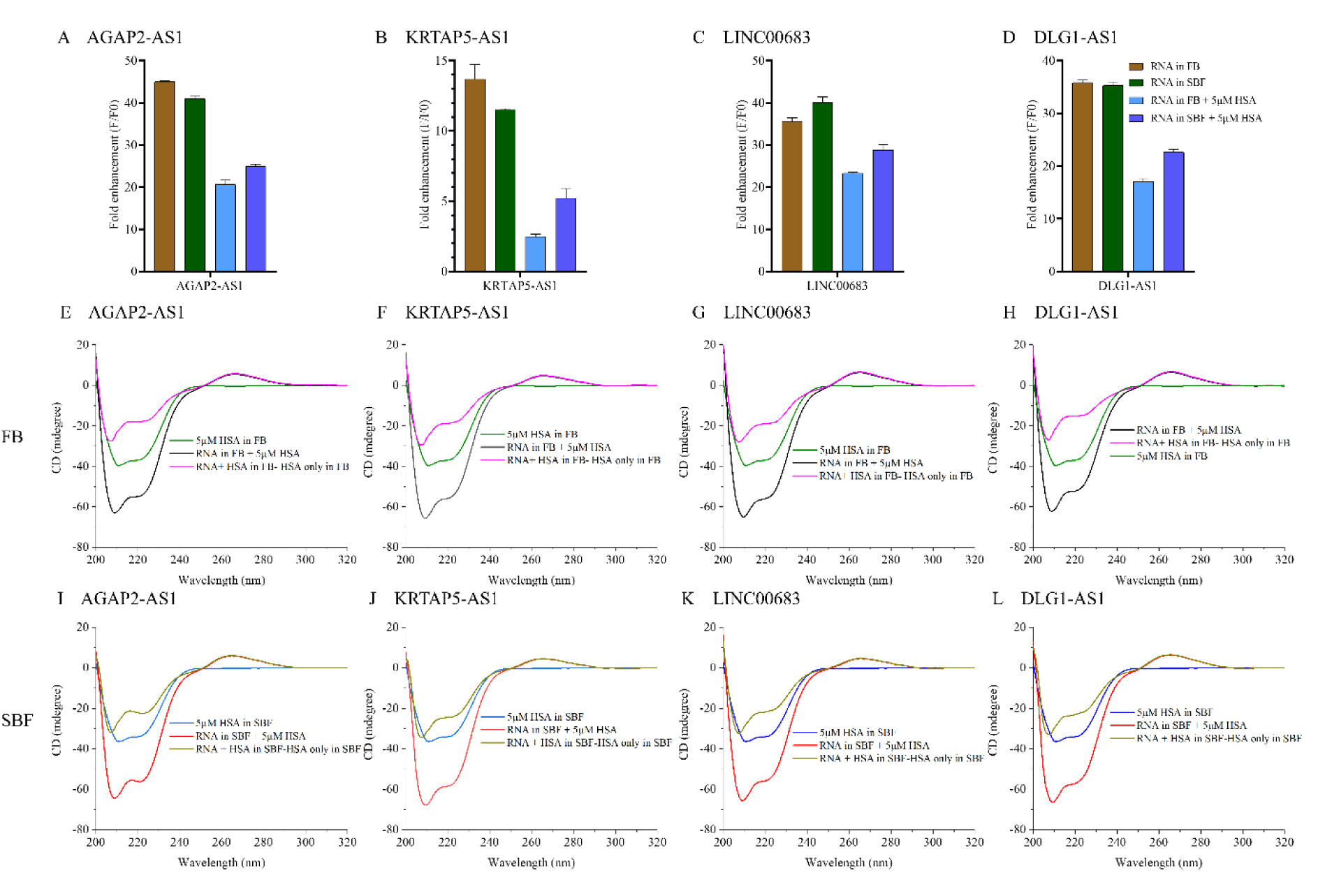
Comparative analysis of PQS structural stability and HSA-induced conformational changes under folding buffer and SBF conditions. (A–D) ThT fluorescence enhancement (F/F0) of AGAP2-AS1, KRTAP5-AS1, LINC00683, and DLG1-AS1 in folding buffer and SBF, with or without 5 μM HSA. (E–H) CD spectra in folding buffer showing HSA alone, RNA + HSA, and corrected spectra generated after subtraction of the HSA-only contribution. (I–L) Corresponding CD spectra in SBF.

While the in silico RPISeq analysis predicted multiple potential interactions for KRTAP5-AS1, our *in vitro* experiments consistently suggested comparatively weaker structural stability and protein interaction behavior. KRTAP5-AS1 exhibited the lowest ThT fluorescence enhancement, weakest HSA binding affinity in ITC analysis, and displayed the sharpest decrease in ThT fluorescence enhancement upon addition of HSA. These observations suggest that while sequence-based algorithms can predict interaction propensity, the actual interaction is strongly influenced by RNA folding, structural accessibility, and conformational stability. Thus, for the purpose of concluding with a more experimentally relevant interaction model, we reconstructed the STRING PPI network after exclusion of proteins that are specifically associated with KRTAP5-AS1 namely HNRPK, ILF3, SFRS1 and FMR1. As shown in **Figure 11**, compared to the original network the refined PPI network appeared more compact and focused, with stronger clustering among RBPs associated with structurally stable G4-forming lncRNAs. Core proteins such as ELAVL1, hnRNPA2B1, IGF2BP1, and FUS remained highly interconnected, whereas ALB, SP1, DHX36 retained a comparatively peripheral position.

**Figure 11.**
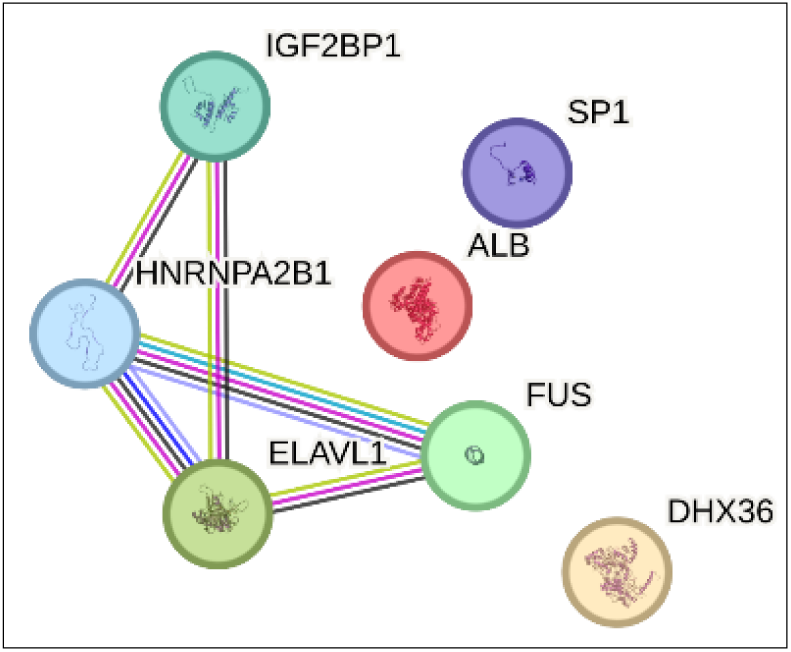
PPI network generated using STRING at a confidence score ≥ 0.7 for RBPs associated with circulating, PQS-containing lncRNAs (excluding KRTAP5-AS1). Nodes represent proteins, with edge thickness indicating interaction confidence and colored edges indicating different sources of supporting evidence. The network contains 7 nodes and 5 edges, with an average node degree of 1.43 and a local clustering coefficient of 0.476. The PPI enrichment p-value (8.74 × 10⁻⁵) indicates statistically significant interaction enrichment.

The proteins IGF2BP1, hnRNPA2B1, and ELAVL1 are all reported as m6A readers that collectively orchestrate post-transcriptional gene regulation in cancer. Their co-convergence on m6A-modified transcripts stabilizing oncogenic mRNAs such as MYC, E2F1, and SRF positions them as cooperative drivers of tumor proliferation, metabolic reprogramming, EMT, and immune microenvironment remodeling *(Wu et al. 2023; Duan et al. 2024b; Cai et al. 2022)* The recurrent co-overexpression of these proteins across diverse malignancies has been associated with poor overall patient survival *(Müller et al. 2019; Liu et al. 2022)*. Their retention within the refined network supports the hypothesis that structurally stable lncRNA G4s preferentially associate with RBPs involved in RNA secondary structure recognition and epitranscriptomic regulation, thereby enriching the network for coordinated post-transcriptional regulators linked to oncogenic progression and chemoresistance *(Yankova et al. 2021)*. Furthermore, filtering out proteins exclusively associated with the experimentally less stable KRTAP5-AS1 further refined the network toward RBPs involved in coordinated oncogenic regulatory pathways.

## Conclusion

The differential expression of circulating lncRNAs between healthy and disease samples has supported their projection as biomarkers. The systematic identification of secondary structure motifs such as G4s within dysregulated circulating lncRNAs is uncommon in literature. In this work, we combined computational prediction and experimental validation to investigate the structural and functional properties of PQS in circulating lncRNAs associated with cancer. Further, through integrative analysis using LncTarD, NPInter, G4IPDB, RPISeq, and STRING, we identified a network of G4-binding RNA-binding proteins potentially associated with AGAP2-AS1, DLG1-AS1, KRTAP5-AS1, and LINC00683, highlighting the relevance of G4-mediated RNA-protein interactions in post-transcriptional regulation. The development of lncRNA-associated PPI networks has mostly relied on identification of specific protein interactors of dysregulated lncRNAs followed by visualization of proteins as a network using STRING. In our work, G4 motifs serve as an entry point for identification of proteins with validated G4-binding activity. Except for the work by Zhang and co-workers that concerned DNA G4s in colorectal cancer serum, no existing report appears to have built a G4-anchored RBP-PPI network for the circulating RNA interactome *(Zhang et al. 2024)*. A detailed experimental scrutiny of the four identified PQS reveals nuances in their *in vitro* structural stability and HSA interacting propensity of the corresponding G4s. AGAP2-AS1, DLG1-AS1, LINC00683, and KRTAP5-AS1 exhibited very strong affinity binding with HSA. Notwithstanding the strong interaction with HSA, KRTAP5-AS1 consistently demonstrated weaker structural stability and lower ThT enhancement. Our experimental findings emphasize that the mere presence of PQSs alone is insufficient to predict functional G4 behavior. We sought to refine the PPI network by exclusion of proteins exclusively linked to the experimentally weakest RNA namely KRTAP5-AS1. Such workflow revision is uncommon considering that lncRNA RBP PPI networks are commonly presented as an endpoint in the analytical workflow. The reconstructed STRING PPI network after excluding KRTAP5-AS1 is a more compact and better-supported 7-node network with a statistically significant PPI enrichment p-value. Overall, the specific circulating lncRNA and interacting proteins described in the work help establish a framework for prioritizing biologically relevant circulating G4-forming lncRNAs and demonstrating the feasibility of developing G4-anchored RBP-PPI.

## Author Contributions

Deepshikha Singh: Conceptualization, Investigation, Methodology, Data curation, Validation, Formal Analysis, Visualization, Writing – original draft, Writing – review & editing, Resources. Arkadeep Ghosh: Investigation, Methodology, Data curation, Formal Analysis, Visualization, Writing – original draft, Writing – review & editing. Shruti Mathur: Investigation, Methodology, Data curation, Formal Analysis, Visualization, Writing – original draft, Writing – review & editing. Sanghati Patra: Investigation, Data curation. Safya Nasir: Investigation, Data curation. Rajesh K Hadiya: Investigation, Data curation. Bhaskar Datta: Conceptualization, Methodology, Validation, Writing – original draft, Writing – review & editing, Funding acquisition, Project administration, Resources, Supervision.

## Acknowledgements

B.D. gratefully acknowledges financial support for this work by Gujarat State Biotechnology Mission (GSBTM) vide project no. GSBTM/JD(R&D)/626/22-23/00006262. All the illustrations are created with Biorender.com.

