## Supporting Information for "Putative G-Quadruplex Structures in Cancer-Dysregulated Circulating lncRNAs and their G4-mediated Identification of Protein Interacting Partners"

* Corresponding author.

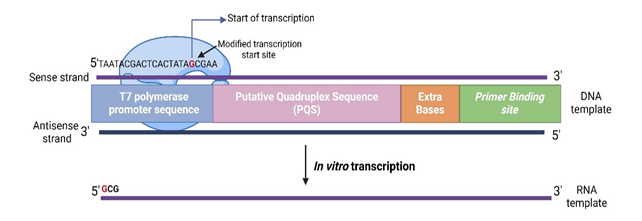

**Figure S1.** Illustration of the design of the DNA templates used for in vitro transcription PQS'. Each template contains a T7 RNA polymerase promoter at the 5′ end, followed by the PQS, a short spacer region, and a primer-binding site for reverse transcriptase stop assay. T7 polymerase initiates transcription at a defined start site to generate RNA transcripts that include the PQS and adjacent regions, enabling downstream assays of G4 folding and RNA–protein interactions.

**Table S1.** Ion concentration used to prepare the SBF

| **Ions** | **Ion concentrations (mm)** |
| --- | --- |
| Na^+^ | 142.0 |
| K^+^ | 5.0 |
| Mg^2+^ | 1.5 |
| Ca^2+^ | 2.5 |
| Cl^−^ | 147.8 |
| HCO_3_^−^ | 4.2 |
| HPO_4_^2−^ | 1.0 |
| SO_4_^2−^ | 0.5 |
| pH | 7.40 |

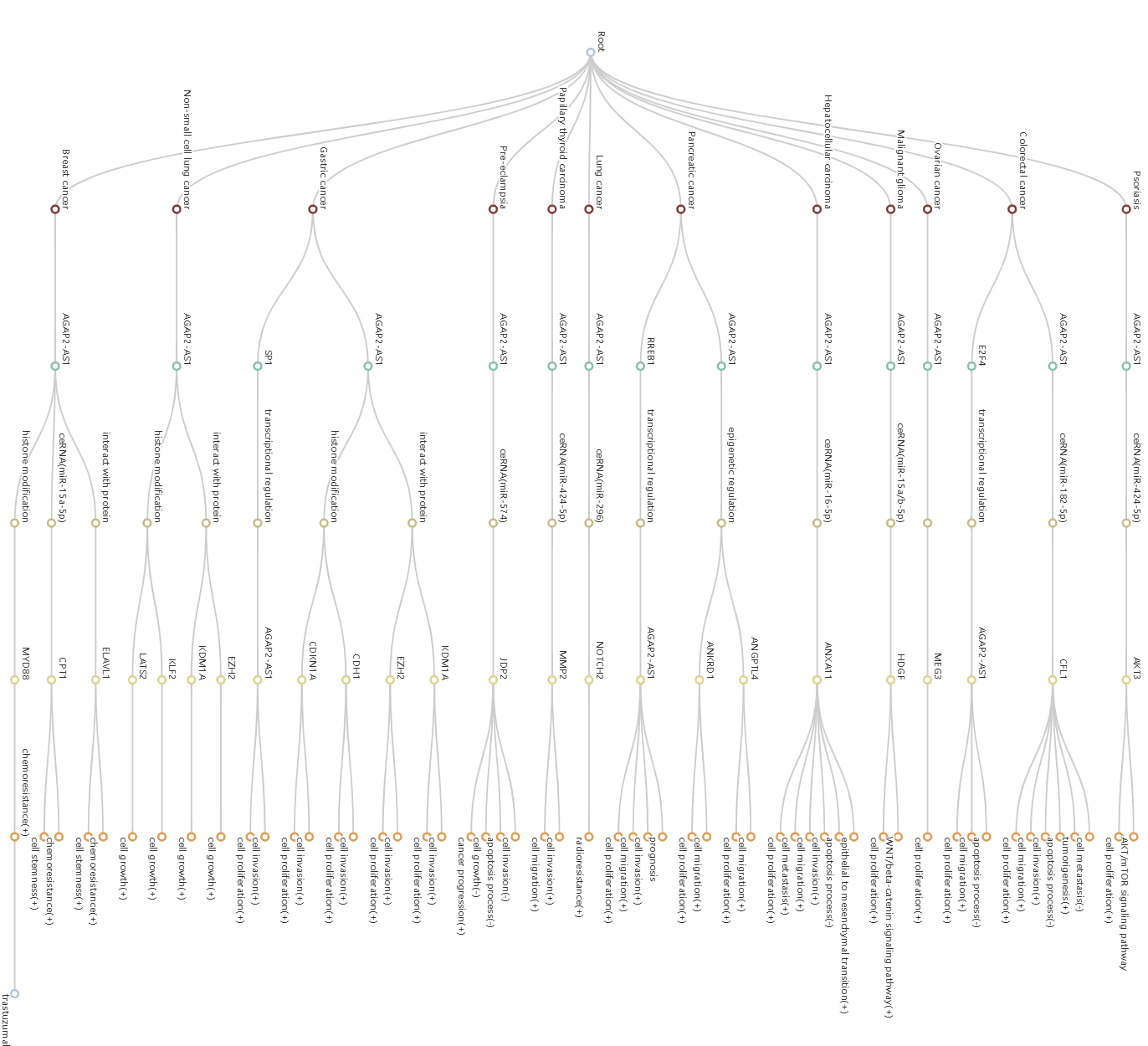

**Figure S2.** Regulatory landscape and interaction network for the lncRNA AGAP2-AS1. Connections link AGAP2-AS1 to diverse cancers and regulatory mechanisms, including ceRNA activity, transcriptional and epigenetic modulation, protein interactions, and pathway engagement. Downstream nodes highlight specific genes and cellular functions affected, such as proliferation, migration, invasion, chemoresistance, and stemness, based on evidence from integrative databases.

**Table S2.** Predicted RPISeq Interaction Scores for Circulating lncRNAs and G4-Binding proteins

| **lncRNA** | **Protein** | **RF Classifier** | **SVM Classifier** |
| --- | --- | --- | --- |
| AGAP2-AS1 | ALB | 0.65 | 0.22 |
|  | DHX36 | 0.65 | 0.95 |
|  | ELAVL1 | 0.65 | 0.32 |
| KRTAP5-AS1 | ALB | 0.80 | 0.23 |
|  | FUS | 0.90 | 0.16 |
|  | hnRNPA2B1 | 0.75 | 0.16 |
|  | SP1 | 0.75 | 0.28 |
|  | DHX36 | 0.75 | 0.89 |
|  | HNRPK | 0.90 | 0.48 |
|  | ILF3 | 0.85 | 0.46 |
|  | SFRS1 | 0.90 | 0.31 |
|  | FMR1 | 0.90 | 0.36 |
| LINC00683 | ALB | 0.85 | 0.79 |
|  | FUS | 0.95 | 0.81 |
|  | IGF2BP1 | 0.85 | 0.84 |
|  | hnRNPA2B1 | 0.70 | 0.65 |
|  | SP1 | 0.80 | 0.81 |
|  | DHX36 | 0.7 | 0.97 |
| DLG1-AS1 | ALB | 0.9 | 0.84 |
|  | FUS | 0.9 | 0.88 |
|  | IGF2BP1 | 0.90 | 0.89 |
|  | hnRNPA2B1 | 0.85 | 0.76 |
|  | SP1 | 0.95 | 0.90 |

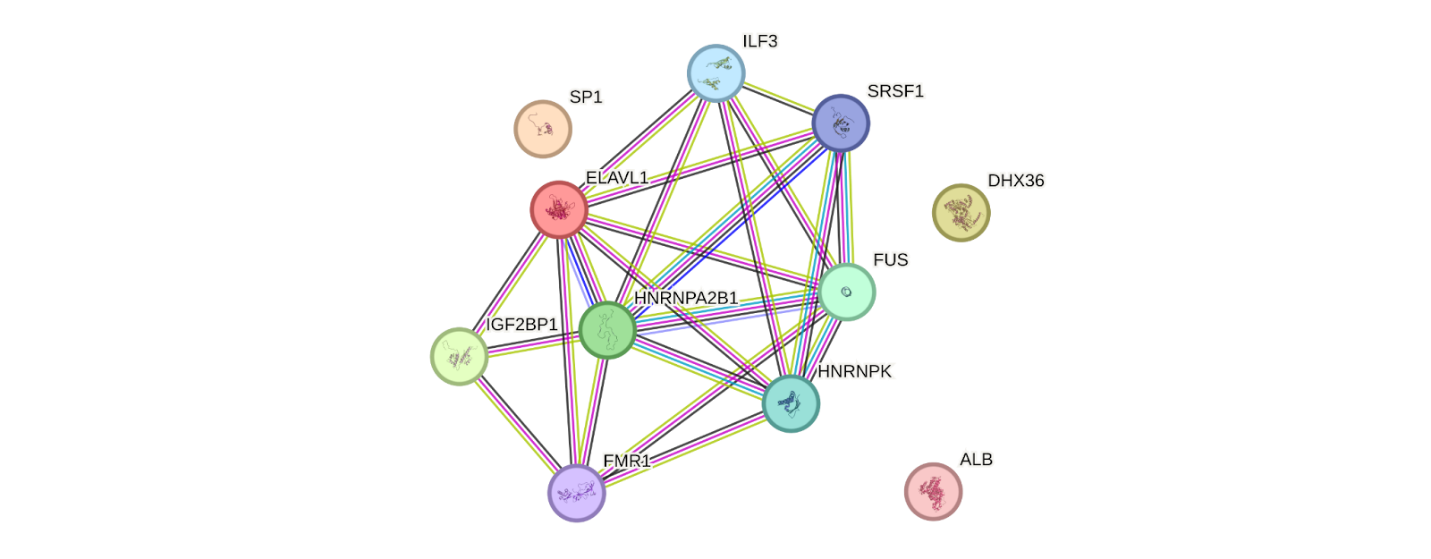

**Figure S3.** High-confidence PPI network (STRING; confidence score ≥ 0.7) of core RNA-binding proteins (RBPs) interacting with dysregulated, PQS-containing lncRNAs. Nodes represent individual RBPs, with edge thickness corresponding to interaction confidence. Central network hubs; ELAVL1, HNRNPA2B1, IGF2BP1, FUS, HNRNPK, SRSF1, ILF3, and FMR1; form a densely interconnected cluster, while DHX36, SP1, and ALB are more peripheral. The significance of network enrichment (PPI enrichment p-value < 1.0e-16) reflects collective involvement in shared molecular functions related to mRNA fate, stability, and post-transcriptional gene regulation.

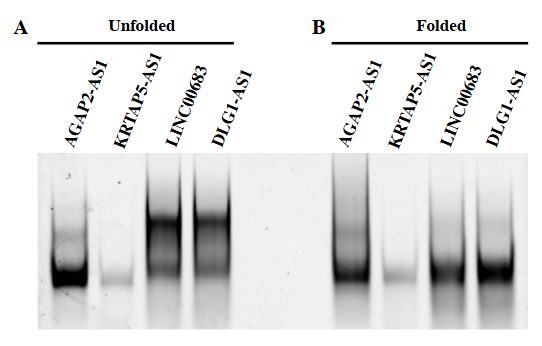

**Figure S4.** Native PAGE gel images ThT staining of AGAP2-AS1, KRTAP5-AS1, LINC00683, and DLG1-AS1. A. PQS' in the unfolded state and B. PQS' in the folded state.

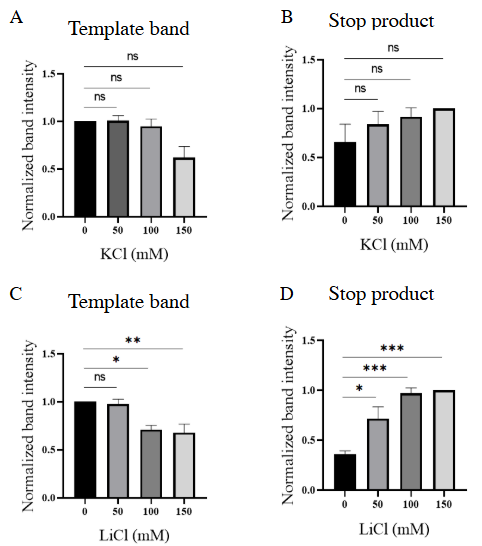

**Figure S5**. RT-stop assays of KRTAP5-AS1, under increasing concentrations of KCl and LiCl. Quantification of (A) and (C) full-length template bands, and (B) and (D) stop product for KRTA5-AS1. Ordinary one-way ANOVA was employed for statistical analysis, and the resulting statistical significance is denoted with asterisks (*). Nonsignificant P-values are represented as ns.

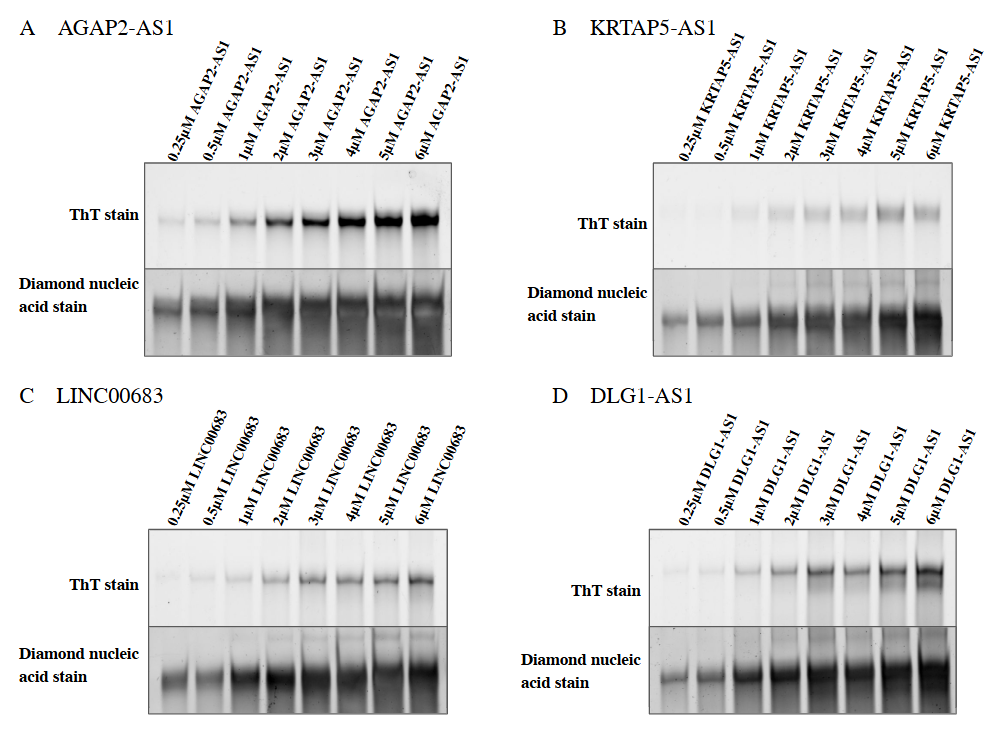

**Figure S6.** Native PAGE gel images of A. AGAP2-AS1, B. KRTAP5-AS1, C. LINC00683, D. DLG1-AS1 in increasing concentration (0.25 μM- 5 μM) visualized after ThT staining followed by diamond nucleic acid staining.

**Table S3.** Thermodynamic parameters of RNA–ligand interactions determined by isothermal titration calorimetry

| PQS of lncRNA | K_d_ | ΔH (Kcal/mol) | ΔG (Kcal/mol) | -TΔS (Kcal/mol) |
| --- | --- | --- | --- | --- |
| **AGAP2-AS1** | 70 nM | -78.1 | -9.76 | 68.3 |
| **KRTAP5-AS1** | 59.3 nM | -80 | -9.86 | 70.1 |
| **LINC00683** | 19.4 nM | -79.9 | -10.5 | 69.4 |
| **DLG1-AS1** | 55.2 nM | -79.9 | -9.91 | 70 |

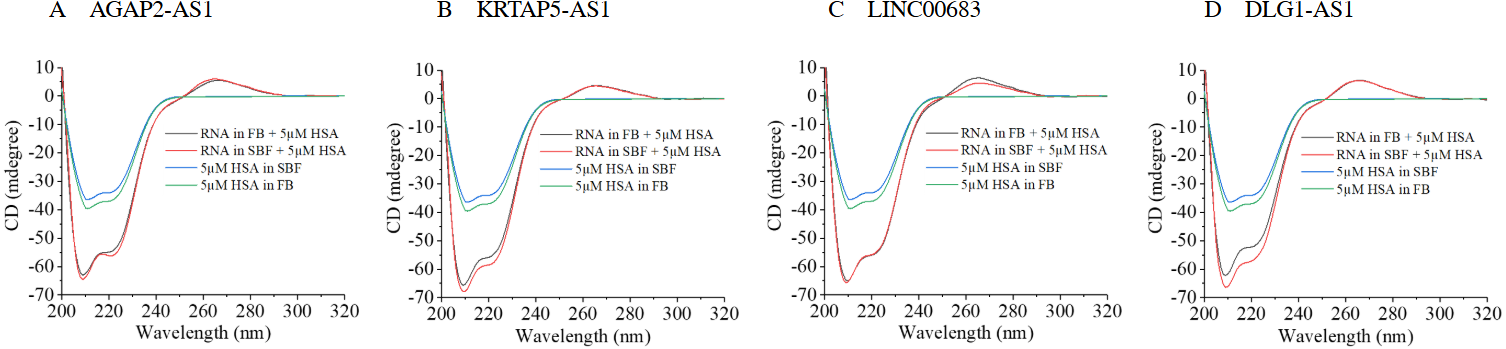

**Figure S7.** Comparative CD of PQS-HSA interactions in folding buffer and SBF. CD spectra of (A) AGAP2-AS1, (B) KRTAP5-AS1, (C) LINC00683, and (D) DLG1-AS1 recorded in the presence of 5 μM HSA under folding buffer and SBF conditions, alongside spectra of HSA alone in the corresponding buffers.

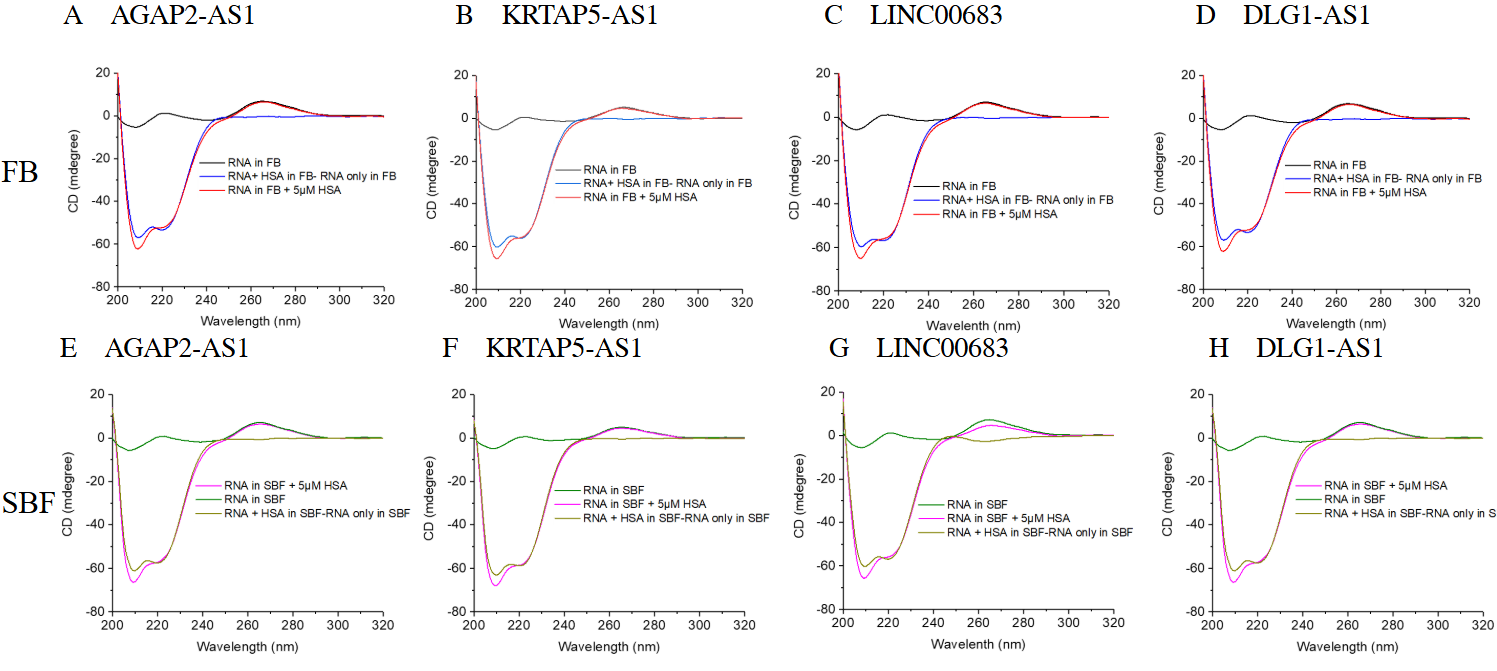

**Figure S8.** Comparative analysis of PQS' structural stability and HSA-induced conformational changes under folding buffer and SBF conditions. (A-D) CD spectra in folding buffer showing RNA alone, RNA + HSA, and corrected spectra generated after subtraction of the RNA-only contribution. (E-H) Corresponding CD spectra in SBF.

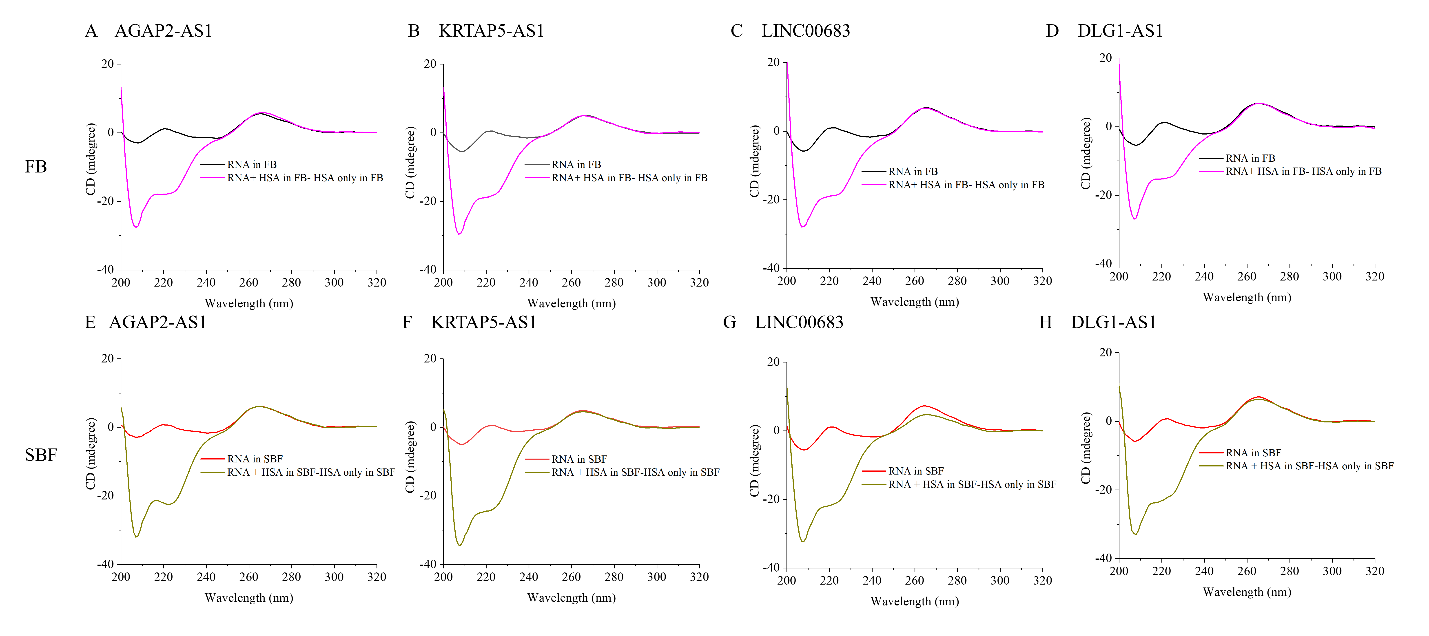

**Figure S9.** Comparative analysis of PQS' structural stability and HSA-induced conformational changes under folding buffer and SBF conditions. CD spectra in folding buffer showing RNA alone, and corrected spectra generated after subtraction of the HSA-only contribution from RNA + HSA spectra. (E-H) Corresponding CD spectra in SBF.
